# Phylum barrier and *Escherichia coli* intra-species phylogeny drive the acquisition of resistome in *E. coli*

**DOI:** 10.1101/2020.10.26.345488

**Authors:** Marie Petitjean, Bénédicte Condamine, Erick Denamur, Etienne Ruppé

**Affiliations:** Université de Paris, IAME, INSERM, F-75018 Paris, France; APHP, Laboratoire de Génétique Moléculaire, Hôpital Bichat, F-75018 Paris, France; APHP, Laboratoire de Bactériologie, Hôpital Bichat, F-75018 Paris, France

**Keywords:** *Escherichia coli*, antibiotic resistance genes

## Abstract

*Escherichia coli* is a ubiquitous bacterium that has widely been exposed to antibiotics over the last 70 years. It has adapted by acquiring different antibiotic resistance genes (ARG), which we aim at characterizing the census here. To do so, we analysed 70,301 *E. coli* genomes obtained from the EnteroBase database and detected 1,027,651 ARG using the AMRFinder, Mustard and ResfinderFG ARG databases. We observed a strong phylogroup/clonal lineage specific distribution of some ARG, arguing for epistasis between ARG and the strain genetic background. However, each phylogroup had ARG conferring a similar resistance pattern, indicating phenotypic convergence. The GC content or the type of ARG was not associated to the frequency of the ARG in the database. Besides, we identified ARG from anaerobic, non-Proteobacteria bacteria in four genomes of *E. coli* supporting that the transfer between anaerobic bacteria and *E. coli* can spontaneously occur but remain exceptional. In conclusion, we showed that phylum barrier and intra-species phylogenetic history are major drivers of the acquisition of resistome in *E. coli*.

## Introduction

*Escherichia coli* is a ubiquitous bacterium found in the intestinal microbiota of vertebrates. In the human gut microbiota, *E. coli* is the dominant species of the phylum Proteobacteria with an average 10^8^ colony-forming units per gram of faeces (Slanetz and Bartley 1957; Tenaillon et al. 2010). It is also commonly found in the digestive tract of mammals including livestock (Smati et al. 2015). Hence during the last 70 years, *E. coli* strains have been highly exposed to antibiotics used in human and animal health as well as those used in agriculture. In response, *E. coli* has adapted by the acquisition of multiple genes encoding antibiotic resistance referred to as antibiotic resistance genes (ARG).

In the pre-next generation sequencing (NGS) era, an exhaustive characterization of ARG in a given species was challenged by the necessity to use as many targeted PCRs than the genes aimed to be detected and perform experiments in a high number of isolates. With the development of NGS and genomics after 2005, the number of bacterial genomes made available has exponentially increased and the identification of ARG has become easier via the use of *in silico* tools. Efforts have been made to collect and organize the known ARG sequences into dedicated databases with the first antibiotic resistance database (ARDB) being released as early as 2008 (Liu and Pop 2009; Zankari et al. 2012). Since then others such as ResFinder (Zankari et al. 2012), CARD (Jia et al. 2017), ARG-ANNOT (Gupta et al. 2014) and more recently AMRFinder (Feldgarden et al. 2019) have followed. Typically, these databases include thousands of ARG nucleotidic and/or amino-acid sequences previously identified in cultivable and/or pathogenic bacteria. However, their content is biased as they lack the ARG found in bacteria that are hard to culture and/or of little relevance from medical perspective such as commensal, strict anaerobic bacteria from the gut microbiota. Nonetheless, we and others have found these bacteria do harbour a vast diversity of ARG, and the latter actually differ from those found in the ARG databases (Ruppé et al. 2019). As for the aforementioned ARG databases, these “other” ARG have been made available into dedicated databases namely FARMEDB (Wallace et al. 2017), ResFinderFG (Munk et al. 2018) and Mustard (Ruppé et al. 2019). Strikingly, the ARG content of the database barely overlapped with that of the other ARG databases. Indeed, very few observations support the transfer of the ARG from intestinal commensals to Proteobacteria opportunistic pathogens such as *E. coli*. Still, such transfer has proven to be possible. For instance, *tet(X)*, a gene encoding resistance to tetracyclines (Deng et al. 2014) that transferred from Bacteroidetes to Proteobacteria. Furthermore, some transfer events may have gone unseen due to the lack of genomic monitoring on a large number of strains.

While most of these ARG are borne by mobile genetic elements such as integrons (Stalder et al. 2012), plasmids (Branger et al. 2018, 2019) or more rarely phages (Billard-Pomares et al. 2014), some associations between their presence and specific *E. coli* phylogroups have been remarked (Johnson et al. 1991, 1994; Branger et al. 2005; Deschamps et al. 2009) as witnessed by the worldwide dissemination of multidrug-resistant clonal groups such as the clonal group A (Manges et al. 2001) and more recently the ST131 (Nicolas-Chanoine et al. 2014). In such clonal groups, strong associations have been evidenced between the within ST sub-clade, the plasmid type and the ARG content (Kondratyeva et al. 2020). All these data argue for a complex cross talk between the chromosomal background, the genetic support of the ARG and the ARG itself resulting from intergenic epistasis (Domingo et al. 2019).

Avalaible since December 2015, EnteroBase (Zhou et al. 2020) is a publicly available database including thousands of genomes from *E. coli* and other species (like *Salmonella*, other *Escherichia* species, *Shigella, Clostridoides, Vibrio, Yersinia, Helicobacter* and *Moraxella*). Beyond genomes, EnteroBase includes (at a variable level of completeness though) metadata linked to the strain itself (name, source, location, laboratory, species, serotype, disease and entry/update date) and to its sequencing process (N50, coverage). Still, no data regarding antibiotic resistance are available. In this study, we leveraged the high number of *E. coli* genomes in the EnteroBase to characterize the acquired ARG in *E. coli* and specifically (i) to test for specific associations between the phylogenetic background of the strains and the presence of ARG and (ii) to evidence ARG transfer between anaerobic bacteria and *E. coli* using a metagenomic database ResFinderFG (Munk et al. 2018) and Mustard (Ruppé et al. 2019).

## Results

### *Distribution of the species and* E. coli *phylogroups among the* E. coli/Shigella *EnteroBase*

During the curation step, 5,144 genomes (6.3% of genomes) were re-assigned to another genus/species. After curation, we identified 70,301 *E. coli,* 221 *Escherichia* clade I, 5 *Escherichia* clade II, 26 *Escherichia* clade III, 17 *Escherichia* clade IV, 102 *Escherichia* clade V (*E. marmotae*), 216 *E. albertii*, 36 *E. fergusonii* and 11,139 *Shigella* and enteroinvasive *E. coli* (EIEC). Among *E. coli,* the phylogroups distributed as follows: B1 (25,265), A (12,469), B2 (12,414), E (11,965), D (4,139), F (2,403) and C (1,646) (Figure 1). The three major STs were ST11 (n=10,059, phylogroup E), ST131 (n=5,143, phylogroup B2) and ST10 (n=3,959, phylogroup A), making up 27% of the *E. coli* in the database. Of note, the ST11 includes the O157:H7 shiga-toxin-producing *E. coli* (STEC) strains (Kim et al. 2017), while the ST131 includes the O25b:H4 extra-intestinal pathogenic (ExPEC) clone (Nicolas-Chanoine et al. 2014).

**Figure 1:**
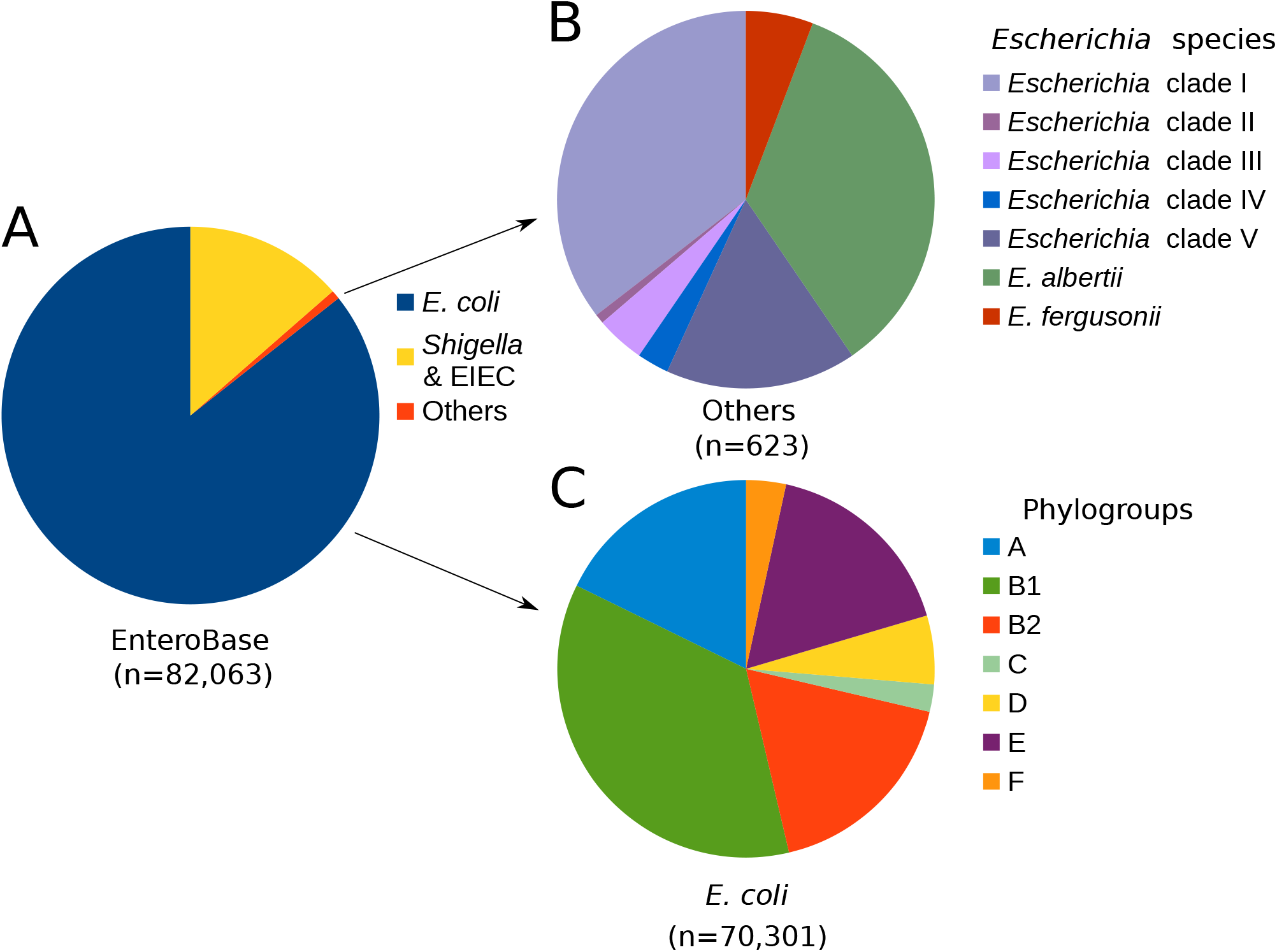
Distribution of species/phylogroups in the *Escherichia/Shigella* EnteroBase. (A) Among the 82,063 genomes of *Escherichia/Shigella* EnteroBase we identified (B) 623 genomes from *E. albertii, E. fergusonii* and *Escherichia* clades (referred to as “others”) and (C) 70,301 genomes of *E. coli* distributed in seven phylogroups. Of note, F phylogroup in the figure includes both F and G phylogroups (Clermont et al. 2019). *Escherichia* clade III and IV are two sub-species belonging to a unique species.

### *The most frequent antibiotic resistance genes found in* E. coli

First, we used the AMRFinder database and identified a total of 314,091 ARG in *E. coli* genomes. The first part (n=164,519) included genes matching to known genes with 100% of identity and coverage. This corresponded to 381 ARG out of the 4,955 genes included in the AMRFinder database. The second part (n=149,572) was made of variants sharing >= 80% of identity and/or 80% of coverage for nucleotide sequences with known genes. This made up variants for 328 genes (including 169 genes not previously detected when the 100% of identity and coverage parameters applied) in AMRFinder database. A total of 550 genes out of 4,955 (11.1%) of the AMRFinder database were thus found at least once in *E. coli* genomes. The 20 most frequent ARG sharing 100% with genes from AMRFinder is depicted in the Figure 2. We predominantly found genes coding for betalactamases and aminoglycosides modifying enzymes (AME), the three most abundant genes being *bla*_TEM-1_ (n=16,766), *aph(3”)-Ib* (n=15,481) and *aph(6)-Id* (n=12,845).

**Figure 2:**
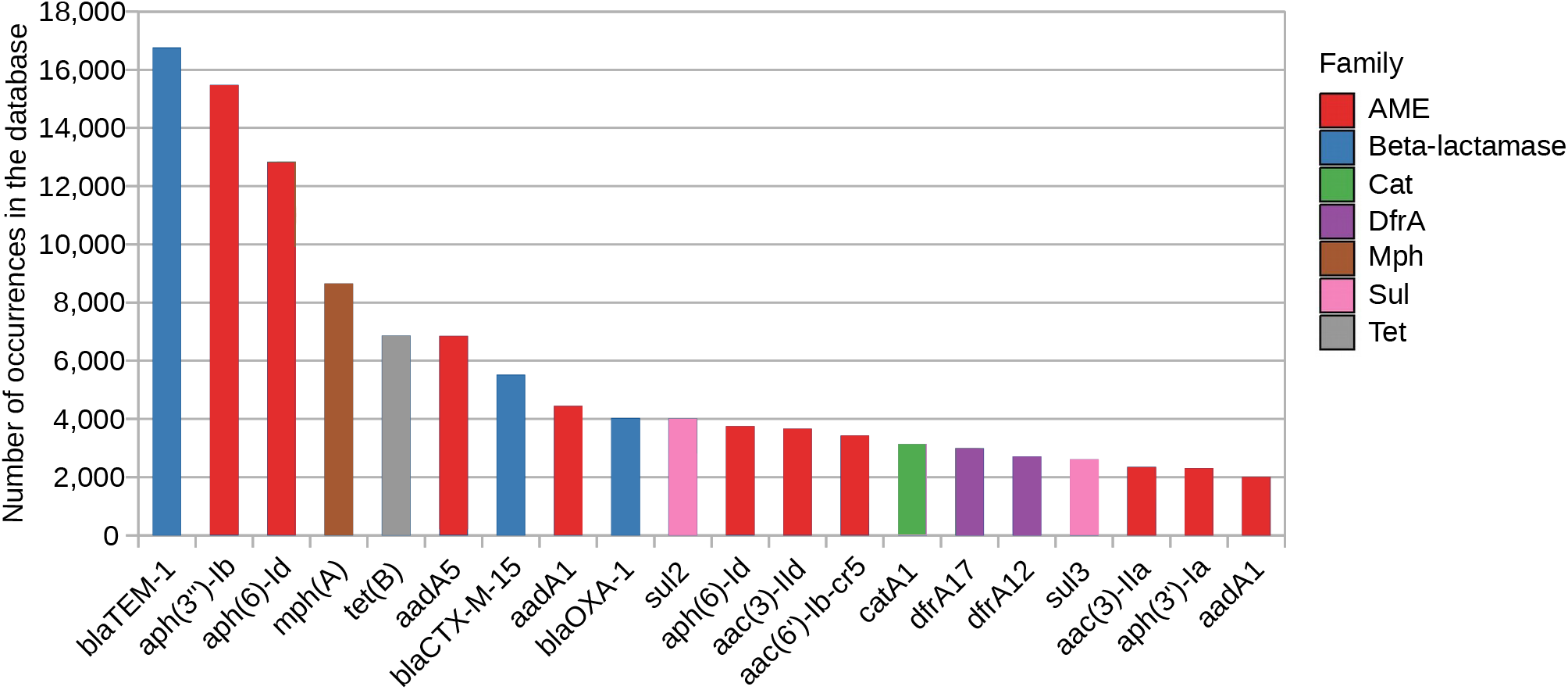
The 20 most frequent antibiotic resistance genes (ARG) mostly detected in the *E. coli* from the EnteroBase database sharing 100 % identity with ARG from AMRFinder. AME: aminoglycoside modifying enzyme; Cat: chloramphenicol acetyltransferase; DfrA: dihydrofolate reductase type A; Mph: macrolide phosphotransferase; Sul: dihydropteroate synthase; Tet: tetracycline efflux pump.

Among beta-lactamases, 27 sub-families were identified (with 100 % identity or in variants) among the 234 found in AMRFinder database. The five beta-lactamases *bla*_TEM_ (n=18,364), *bla*_CTX-M_ (n=10,640), *bla*_OXA-1-like_ (n=4,075), *bla*_CMY-2-like_ (n=1,994) and *bla*_SHV_ (n=518) were the most frequent (Supplementary Table 1).

We also observed a high frequency of ARG conferring resistance to antibiotics that are used to treat infections not caused by *E. coli* but rather caused by Gram-positive bacteria such as rifampicin (*arr*, n=394/110 respectively 100% identity and variants) and macrolide-lincosamide (*lnu* [n=494/322], *mef* [n=258/21], *mphA* [n=8,661/408], *erm* [n=81/1,405], *vga* [n=1/0] and *msr* [n=76/2]).

Unexpectedly, we identified a *blaZ* gene commonly found in *Staphylococcus aureus* in an *E. coli* strain. However, a subsequent analysis of the genome revealed that 10% of reads were assigned to *S. aureus* and 90% to *E. coli*. This 10% of reads from *S. aureus* evenly distributed in the genome of *S. aureus* strain CFSAN007851, strongly supporting a contamination prior to sequencing.

### Distribution of the resistance genes according to the strain phylogeny

After normalization, we observed a variable distribution of the ARG richness in each phylogroup with respectively 254, 236, 213, 170, 178, 122 and 169 genes in phylogroups A, B1, B2, C, D, E and F. However, we observed an even distribution of the diversity with Shannon index respectively equal to 3.63, 3.51, 3.15, 3.58, 3.40, 2.92 and 3.51. The three major genes in each phylogroup were *bla*_TEM-1_, *aph(3”)-Ib* and *aph(6)-Id*, except for phylogroup D and F in which, respectively *mph(A)* (a phosphotransferase conferring resistance to macrolides) and *tet(B)* (an efflux pump conferring resistance to tetracyclines) ranked second and third. However, specific ARG were more frequently found in some phylogroups (referred as phylogroup-predominant resistance gene PPRG, *i.e*. ARG with a p-value less than 0.001 with Kruskal-Wallis test and Benjamini-Hochberg correction, Figure 3). Indeed, 53.5% (n=197/368) of the *bla*_OXA-48_ (a widely spread carbapenemase-encoding gene) were found in phylogroup D. Likewise, 49.5% (n=4,141/8,366) of *bla*_CTX-M-15_ (encoding the worldwide most prevalent extended-spectrum beta-lactamase) were found in phylogroups B2 and C (25% each). Besides, phylogroups B1 and E represented 25,265 and 11,965 genomes but only 5 and 35 PPRG, respectively. In contrast, we observed 102 PPRGs in the phylogroup A (n=12,469).

**Figure 3:**
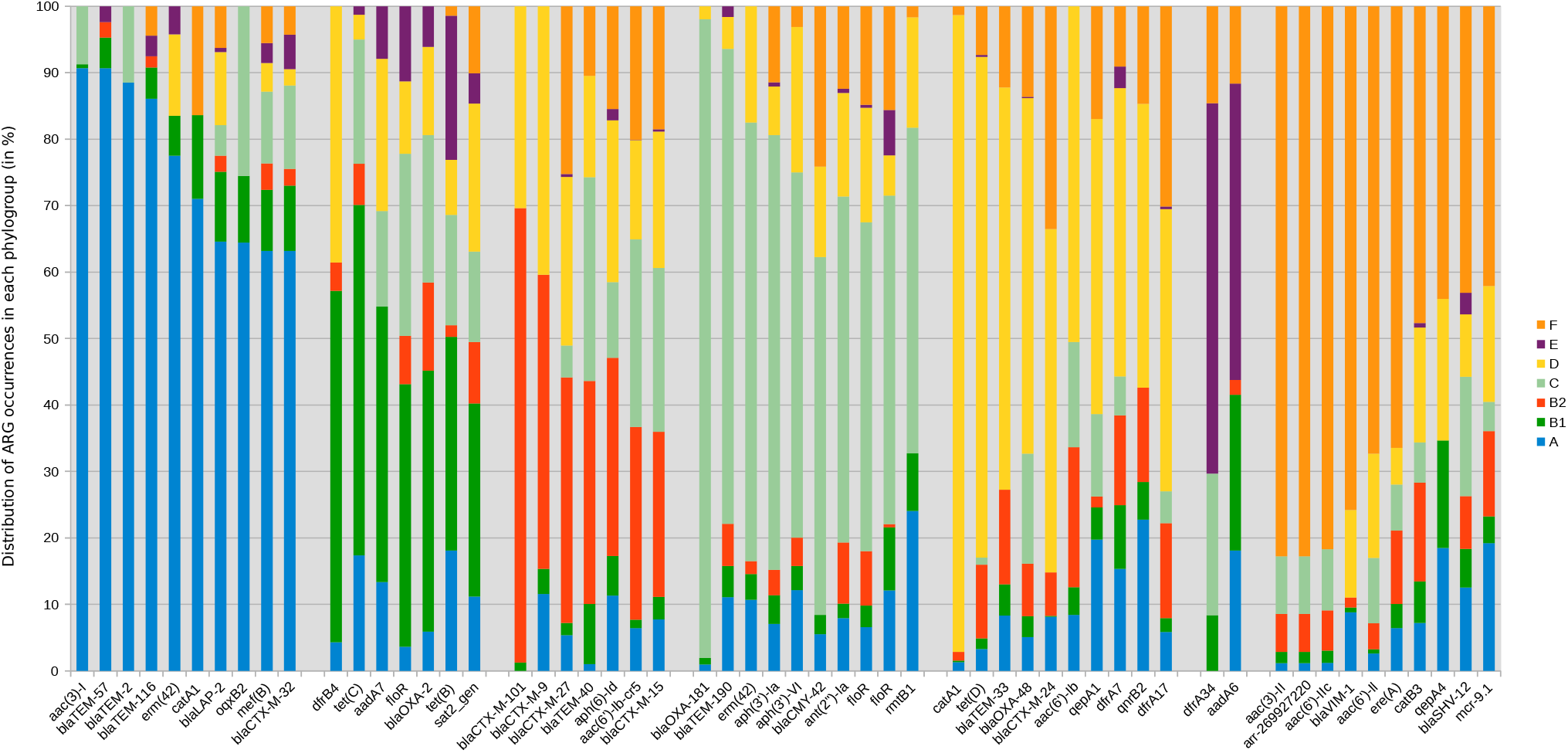
Histogram of the 10 (when available) most representative antibiotic resistance gene in each *E. coli* phylogroup. Only genes with at least 20 occurrences in the database were represented. For each phylogroup, genes were sorted by decreasing frequencies.

Then, we tested the hypothesis that even if the distribution of ARG differed according of the phylogroup, that of their function (*i.e*. the antibiotic families they encode resistance to) would not. We applied the same statistical approach and indeed, found no specific association between the spectrum of ARG families and phylogroups (Figure 4).

**Figure 4:**
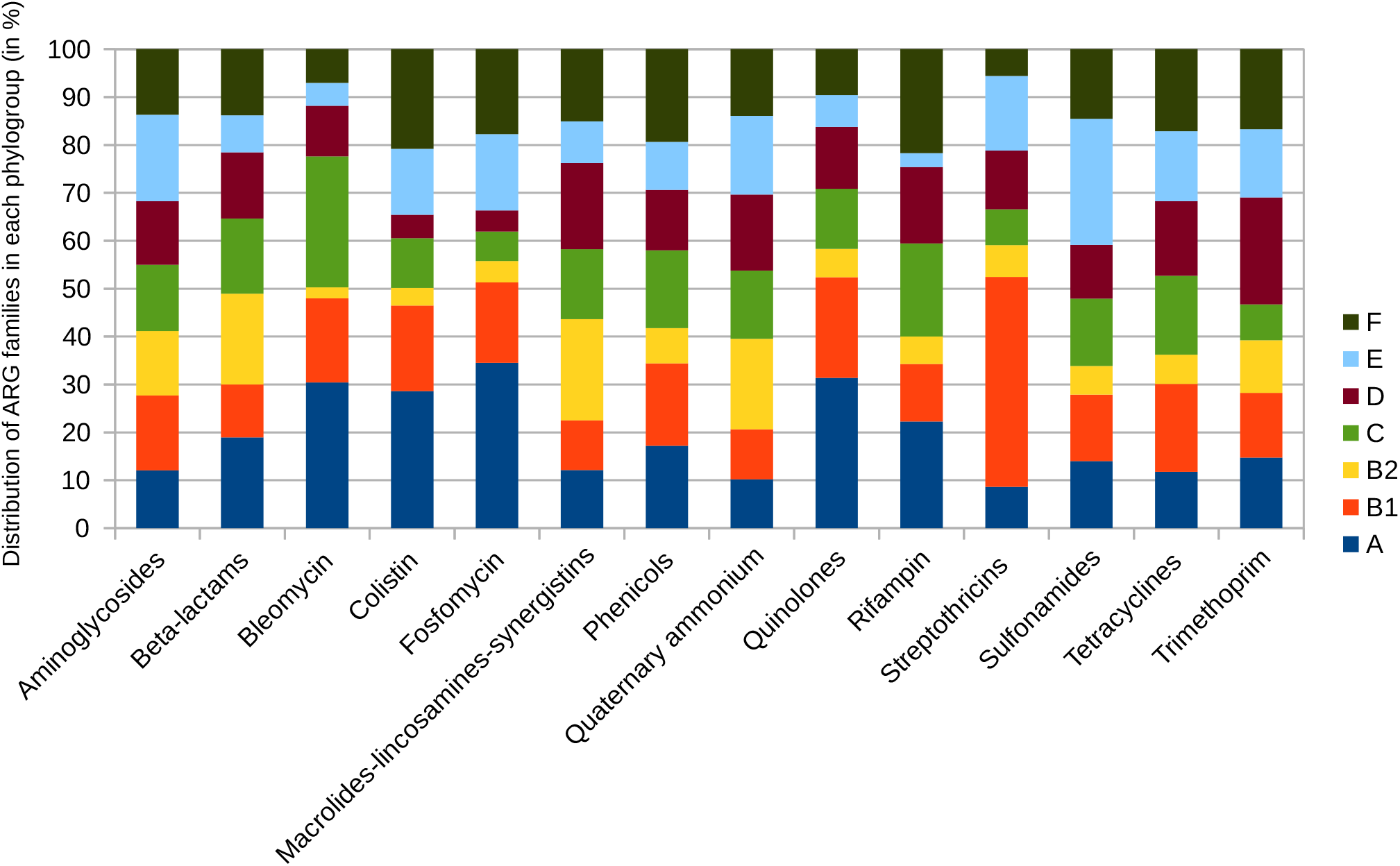
Histogram of the distribution of the targeted antibiotic families in each *E. coli* phylogroup.

**Figure 5:**
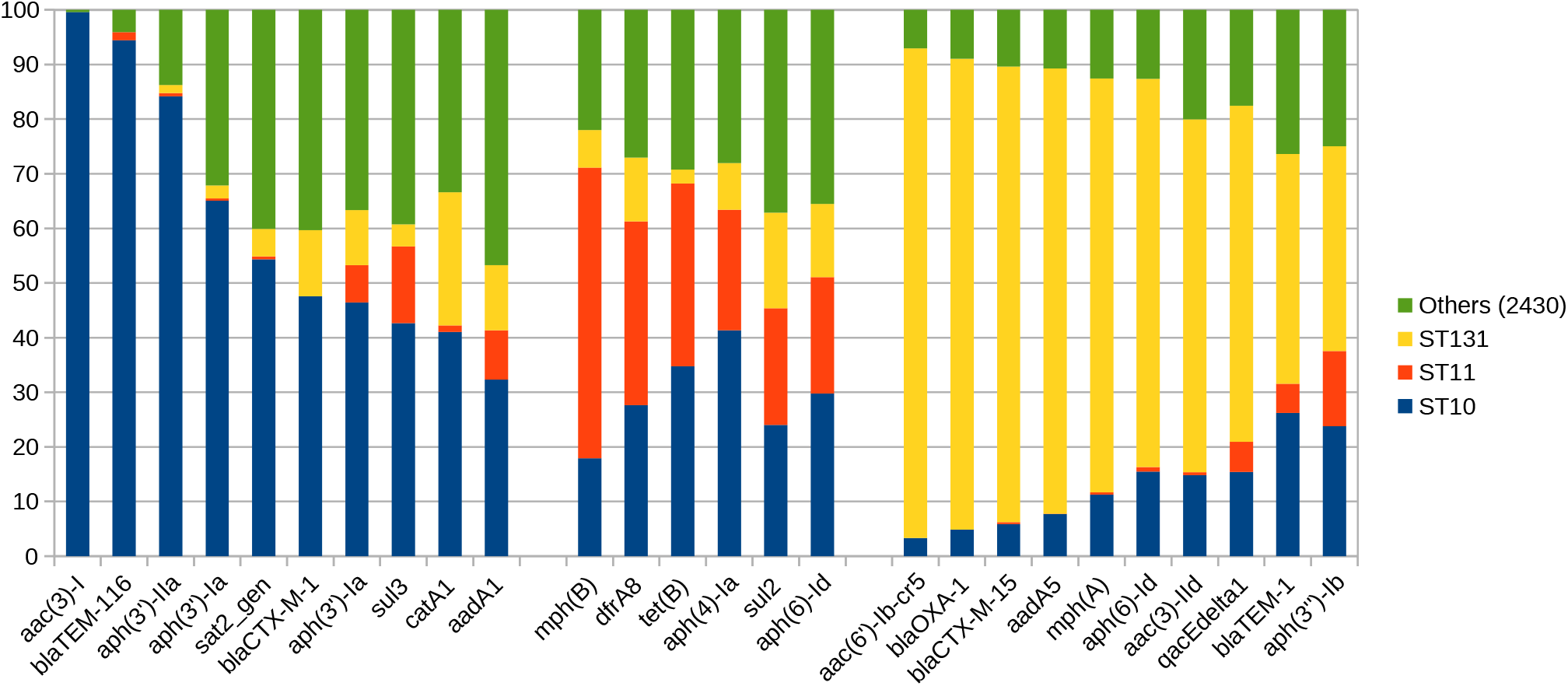
Histogram of the 10 (if available) most representative ARG in each *E. coli* major ST. Only genes with at least 20 occurrences in the database were represented. For each ST, genes were sorted by decreasing frequencies. Others representing all the 2,430 others STs with few occurrences of the indicating genes. Each gene was preferentially found in one ST with a p-value less than 0.01 with Kruskal-Wallis test and Benjamini-Hochberg correction.

We then searched for differential abundances with respect to the main STs of the EnteroBase (ST11 [n=9,391], ST131 [n=4,841] and ST10 [n=3,407]). Despite ST11 had the maximum number of genomes in the EnteroBase, its ARG richness was 65 while it was 162 and 169 in ST131 and ST10, respectively. The Shannon entropy for ST11, ST131 and ST10 were 2.36, 2.86 and 3.12, respectively. We also sought for a specific distribution of some widely spread, clinically relevant beta-lactamases. *bla*_TEM-1_ and *bla*_CTX-M-15_ were mostly found in ST131 (12.5% [n=2,094] and 42% [n=2,310]). Conversely, *bla*_NDM-1_ was mostly found in the ST101 (18%, n=38).

### Global associations between strain phylogeny, plasmid type and resistance genes

We tested correlations between ARG, phylogroups, STs and plasmid incompatibility groups. A total of 124 clusters were found with a correlation factor strictly higher than 0.30, each containing at least one ARG with other ARG, STs, phylogroups or plasmid incompatibility groups. The *bla*_CTX-M-15_ gene strongly correlated with the *aac(6’)-Ib* and *bla*_OXA-1_ (r=0.70) and also in a lesser extent correlated to *aac(3)-IIa, mphA, aadA5, qacEdelta1* and the ST131 as well as the plasmid incompatibility group IncFII (r=0.36). In contrast, we did not identify other ARG, plasmid incompatibility group or ST associated to the other common *bla*_CTX-M_ genes *bla*_CTX-M-27_ and *bla*_CTX-M-1_. As for carbapenemase-encoding genes, a correlation was detected between *bla*_NDM-1_ and *aph(3’)-VI, floR, erm-42, bla_CMY-6_, mphE, msr(E), armA* and with the plasmid incompatibility group IncA (r=0.36). Last, *bla*_TEM-1_ correlated with *aph(3”)-Ib, aph(6)-Id, aac(3)-IId, sul2* and the incompatibility group IncQ1 (r=0.39) in coherence with them being found on the same contig in 97% instances (n=16,263/16,766). Of note, we did not observe any negative correlation between ARG. However, the genes *tet(B), aadA5, qacEdelta1, bla_TEM-1_, bla_CTX-M-15_, bla_OXA-1_* and *mph(A)* were negatively correlated to the ST11.

### GC content and type of antibiotic resistance

We determined the GC content of clusters of ARG and the distribution of their divergence with the mean GC of *E. coli* as measured from the EnteroBase genomes. We observed a large panel of GC content deviation (Figure 6) between the mean GC content of *E. coli* core genome (Bohlin et al. 2017) and the GC content of acquired ARG, supporting that the GC content was not a constraint for the acquisition of ARG from other species or within the *E. coli* species. Interestingly, we also observed that the GC of the most frequent ARG found in *E. coli* (with at least 1,000 occurrences) did not significantly differ from that of the ARG with low frequency (Mann-Whitney test, p=0.7, mean most frequent ARG 50.0 and 48.6 for low frequency).

**Figure 6:**
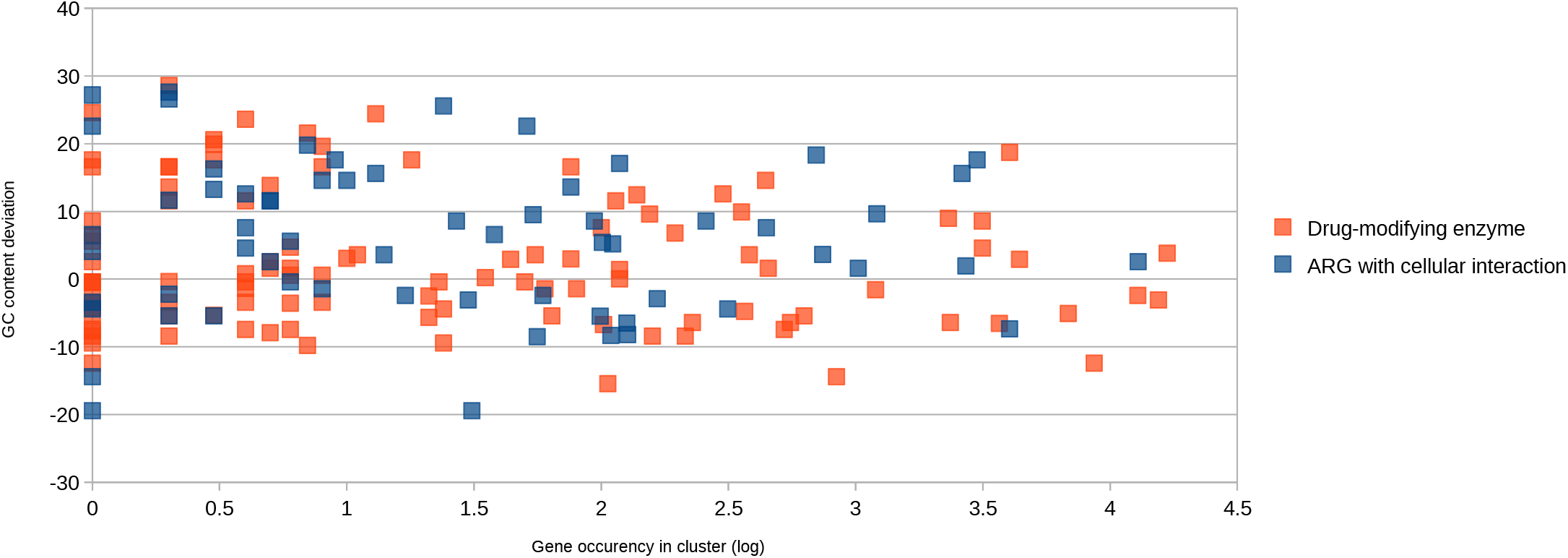
Scatter plot of the GC deviation of ARG according to their occurrence in EnteroBase and the type of resistance encoded. Each dot corresponds to one cluster of ARG (90% nucleotide identity) created using AMRFinder, Mustard and ResfinderFG databases. The GC content deviation corresponds to the difference between the mean GC content of *E. coli* (Bohlin et al. 2017) and the mean of the GC content of each gene in the cluster.

The functional class of the horizontally transferred ARG has been shown to play a role on the fitness of the recipient strain in that genes encoding for drug modification (*e.g*. beta-lactamase and AME) would have a minimal impact on the fitness of the recipient strain even if originating from a GC-divergent background. Conversely, genes encoding for proteins interacting with the cellular content (*e.g*. efflux and target modification) would impact the fitness of the recipient, especially when originating from GC-divergent background (Porse et al. 2018). However, in our dataset we did not observe a distinct pattern according to the type of resistance (Mann-Whitney test, p=0.7, mean for drug-modifying enzyme equal to 3.969 and 2.582 for ARG with cellular interactions, Figure 6).

### *Non-proteobacteria antibiotic resistance genes can be exceptionally found in* E. coli

Last, we assessed whether some ARG found in non-Proteobacteria bacteria were present in the *E. coli* genomes of the EnteroBase using the specific databases ResFinderFG and Mustard. We observed a high number of hits (n=833,814) in the Mustard and ResFinderFG databases mostly corresponding to genes predicted to originate from Proteobacteria. The first part (n=271,458) included genes matching to known genes with 100% of identity and coverage. This corresponded to 37 different genes identified in the 6,095 genes of the Mustard database (Supplementary Table 3) and 44 different genes identified in the 2,282 genes of the ResFinderFG database (Supplementary Table 3). The second part (n=562,356) was made of variants sharing >= 80% of identity and/or 80% of coverage for nucleotide sequences with known genes. This made up 385,673 variants for 23 additional genes in Mustard (Supplementary Table 3) and 176,683 variants for 61 additional genes in ResFinderFG (Supplementary Table 3).

Interestingly, we identified a putative beta-lactamase encoding gene from ResFinderFG (beta_lactamase|KU546399.1|feces|AMx) and Mustard (MC3.MG60.AS1.GP1.C4251.G1) database also found in the strain *Bacteroides uniformis* NBRC 113350 (NCBI accession number NZ_AP019724.1). The *E. coli* bearing this gene was from phylogroup A, ST744 (Table 1) and had been isolated in Germany (Mellmann et al. 2016) from a patient screened for multidrug-resistant bacteria at the University Hospital of Münster. The gene was embedded in a 7,600 bp contig (Supplementary Figure 1), itself sharing 100% identity with the *B. uniformis* genome. The strain was kindly provided by Prof. Alexander Mellmann from University of Münster and was re-sequenced in our laboratory using Illumina (San Diego, CA) MiniSeq and Oxford Nanopore (Oxford Nanopore Technologies, UK) chemistries, which confirmed the presence of the resistance gene (data not shown). Further description of this strain is underway and will be detailed in a separate work.

**Table 1:** Description of the antibiotic resistance genes shared between *Escherichia coli* and non-Proteobacteria strains.

| *E. coli* strain ID | Phylogroup | Sequence type (Warwick University/<br>Pasteur Institute schemes) | Serotype | *fimH*<br>allele | Resistance<br>gene | Strain of origin ID |
| --- | --- | --- | --- | --- | --- | --- |
| ESC_HA9845AA | A | 744/2 | Onovel32:H10 | 54 | putative beta-lactamase | *Bacteroides uniformis*<br>strain NBRC 113350 |
| ESC_OA1280AA | E | 753/920 | O130:H9 | 124 | *erm*(49) | *Bifidobacterium breve*<br>strain CECT7263 |
| ESC_JA0734AA | D | 405/477 | O102:H6 | 27 | *erm* | *Clostridioides difficile*<br>strain CDT4 |
| ESC_FA9928AA | A | 5943/999 | O89:H11 | 41 | *tet*(M) | *Clostridioides difficile*<br>strain CD161 |

We also found two *erm* genes (ribosomal methylases), encoding for macrolide-lincosamide resistance and originating from non-Proteobacteria. The first one was found in an *E. coli* from phylogroup E, ST753 originating from a livestock sample from Ireland (Table 1). A part of the 12,910 bp contig containing the gene, matched against a *Bifidobacterium breve* genome (Supplementary Figure 2). We could identify some transposase genes in the contig bearing the *erm* gene and surrounding inverted repeat sequences. The second *erm* was found in an *E. coli* from phylogroup D, ST405 sample from the Netherlands (Table 1) and was originally identified on a plasmid from *Clostridioides difficile* (Supplementary Figure 3). Unfortunately, the metadata associated to those genomes was not sufficient to trace back the strains and confirm the presence of the genes in *E. coli* by resequencing the strains. However, no evidence of wet lab or dry lab contamination was observed.

We identified a *typA* gene (tyrosine phosphorylated protein A) commonly found in *Streptococcus pneumoniae* in an *E. coli* strain. However, the close analysis of the genome revealed that the reads of *S. pneumoniae* D39 sequenced in the same project than the *E. coli* strain covered all the genome of *E. coli*, strongly supporting a contamination.

Last, we identified a *tetM* gene, encoding for tetracycline resistance, originating from *Clostridioides difficile* (Supplementary Figure 4). The gene was found in a 3,291 bp contig completely matched against the *C. difficile* strain in an *E. coli* from phylogroup A, ST5943, originating from Thailand. No evidence of contamination was observed even if the metadata associated to these genome was not sufficient to confirm the presence of the *C. difficile* gene.

## Discussion

Using a large number of genomes, we could assess the diversity and the distribution of ARG in *E. coli*. From a global perspective, we assumed that the richness of acquired ARG in *E. coli* was somewhat limited with regards to the high number ARG in the literature (the ARG found in *E. coli* making 11.1% of the AMRFinder database). This suggested constraints in the pathways for ARG exchanges and sustainability between species (phylogenetic origin of the gene, interaction within the cell of the gene product (Porse et al. 2018; Jain et al. 1999)), but also within the *E. coli* species. We did not evidence any link between the GC content or the functional class of the transferred genes and their frequency in the database. However, some ARG had specific associations with genomic background (phylogroups and STs), other ARG and plasmid incompatibility groups. Of note, we did not observe any negative correlation between ARG, suggesting that the ARG are not competing with each other. Similar association between the genomic background and the presence of virulence genes has been reported within the *E. coli* species (Escobar-Páramo et al. 2004), indicating that epistasis between different parts of the genome (Domingo et al. 2019) is at play for genes important for adaptation. Nonetheless at a wider scale, we observed that these preferential genetic pathways led to the same functional patterns. While *E. coli* phylogroups had different ARG distributions, they harboured a full genetic armamentarium to resist to the same antibiotic families. In brief, the ARG were different but their functions were similar. Such a functional redundancy suggests that the *E. coli* phylogroups were exposed to the same antibiotics but took different genetic pathways to cope with them.

The antibiotic exposure does not only exert on the bacteria hold responsible for an infection but also the bacteria residing in our microbiota. Indeed, we observed a high rate of ARG conferring resistance to antibiotics that are used to treat infections not caused by *E. coli* or other Gram-negative bacteria but rather caused by Grampositive bacteria such as rifampicin and macrolide-lincosamides. Such antibiotics are excreted in the intestine at high concentrations so that some bacteria with minimal inhibitory concentrations (MIC) too high to be in the clinical spectrum of the antibiotic would be affected in the gut. In that, the high frequency of ARG conferring resistance to those antibiotics is a strong signal stressing the impact of antibiotics on our microbiota.

We observed four putative transfers of ARG between non-Proteobacteria and *E. coli*. In a previous work (Sommer et al. 2009), we observed that the vast majority of ARG found in the intestinal microbiota were very distinct from that found in cultivable bacteria (including *E. coli*) and that little arguments were found to support their mobility. Taking advantage of the largest *E. coli* database to date, we could observe that ARG could actually be exchanged between *E. coli* and intestinal commensals, such as it was observed for *tet(X)* (Leski et al. 2013). Even if exceptional in this dataset with four observations, the fact itself that they were detected suggest that they are not that uncommon, but unlike *tet(X)* which has met success, their spread was very limited to date as they were only found in one genome each. The donor bacteria were strict anaerobic bacteria (*B. uniformis, B. breve* and *C. difficile*) commonly found in the gut microbiota at high abundances (Li et al. 2014) aside *E. coli*. Considering that humans have been using antibiotics for more than 70 years, very favourable conditions were met for ARG transfers between anaerobic bacteria and *E. coli*. That such transfers have been observed so rarely support that anaerobic bacteria may indeed provide ARG to *E. coli*, but that its contribution to the worldwide AMR issue seems to be anecdotal. Indeed, the most successful ARG found in *E. coli* originate from other Proteobacteria species *e.g. bla*_CTX-M_ progenitors are *Kluyvera* spp belonging to the Enterobacterales (Bonnet 2004). Besides, the observations of the ARG transfer would not have been possible if not for the use multiple databases to cover the broadest range of ARG. While databases such as AMRFinder may be found suitable to identify ARG from clinically relevant bacteria, they may not be appropriate when it comes to look for ARG originating from other environments such as the gut microbiota.

We acknowledge some limitation. Despite including a large number of genomes, the EnteroBase suffers from inclusion biases in that strains of interest (*e.g*. the resistant and/or the pathogenic ones) are the most sequenced. Indeed, the EnteroBase includes a large number of STEC mainly the O157:H7 serotype, ExPEC with the emerging ST131 and many strains producing extended spectrum beta-lactamases. Hence, we assume we did not cover the global picture of the acquired ARG ever found in *E. coli*, but only a snapshot. Many more ARG may have been found in *E. coli*, perhaps including some from intestinal commensals which were not found of interest to be cultured and sequenced. Nonetheless, we believe our findings are sound with regards to the very high number of strains included in this study.

In all, we observed that ARG distributed in the *E. coli* phylogroups/STs with a preferential fashion, but that eventually they provided resistance to the same antibiotic families. Furthermore, we observed that the transfer of ARG between non-Proteobacteria and *E. coli* indeed occurred but seemed to be exceptional.

## Material and methods

### *Genomic database, species classification and* E. coli *phylogroup determination*

A total of 82,063 available genomes was downloaded from *Escherichia/Shigella* EnteroBase (as of 1^st^ February 2019). The genomes were classified according to their genera and species (*Shigella, E. coli, E. albertii, E. fergusonii, E. marmotae* or unknown). Using the ClermonTyper (Beghain et al. 2018), a tool that provides information about phylogroups (A, B1, B2, C, D, E, F and G) for *E. coli* and identifies nearest species in conjunction with Mash (Ondov et al. 2016) (*E. fergusonii, E. albertii* and *Escherichia* clades), all the genomes were classified as *E. coli* belonging to the aforementioned phylogroups, *E. fergusonii, E. albertii* and *Escherichia* clades I to V. Of note, in this work we considered that phylogroup F included both F and G phylogroups (Clermont et al. 2019). When a discrepancy was observed between the ClermonTyper and Mash attribution (n=16,132 cases) the strain was classified according to Mash. *Shigella* and EIEC genomes were identified using the *in silico* PCR with primers of the *ipaH3* gene (Sahl et al. 2015). Due to their specific, obligatory intracellular pathogenic trait, they were not considered further.

### Antibiotic resistance and virulence gene identification, plasmid incompatibility group, chromosomal multilocus sequence type determination and GC content

Diamond (Buchfink et al. 2015) was used to identify all the ARG in the EnteroBase by aligning all genomes against the AMRFinder (Feldgarden et al. 2019) (2019-04-29 version), Mustard (Ruppé et al. 2019) (2017-09-30 version) and the ResFinderFG (Munk et al. 2018) (2016-12-21 version) databases (80% of identity and/or 80% of coverage minimum for nucleotide sequences). All redundancies between the databases were removed. The ARG families were picked according to the Mustard website (mgps.eu/Mustard). All ResfinderFG and Mustard ARG originated from non proteobacteria were further investigated. Virulence genes were determined using VirulenceFinder (Joensen et al. 2014) with 90% of identity and 80% of coverage. PlasmidFinder (Carattoli et al. 2014) database was used by Diamond (Buchfink et al. 2015) to determine the plasmid incompatibility groups found in each genome of the EnteroBase database (98% of identity and 95% of coverage minimum). Chromosomal multilocus sequence type (ST) determination according to the Warwick University and/or Pasteur Institute was performed by the mlst software (https://github.com/tseemann/mlst) using the pubMLST database (Jolley and Maiden 2010). GC content (%) deviation between *E. coli* and the acquired ARG was measured using the *E. coli* GC defined previously (Bohlin et al. 2017). ARG family was obtained by clustering all the ARG from the three databases (AMRFinder, Mustard and ResfinderFG) using the homology between these genes (90% of identity) and CD-HIT-EST (Fu et al. 2012; Li and Godzik 2006).

### Statistical analysis and normalization

To circumvent the sequencing biases, the data were normalized so that each phylogroup would include the same number of genomes (n=10,000) by up-sizing (for C, D and F phylogroups) or down-sizing (for A, B1, B2 and E phylogroups). Then, we tested the correlations between phylogroup, ST, plasmid incompatibility group and ARG using the *corrplot* package of R v3.4.2 and the *corrmat* function. The preferential distribution of some ARG within phylogroups was tested using Kruskal-Wallis test and Benjamini-Hochberg correction. The diversity of the ARG in some STs was measured using the Shannon index in R (v3.4.2) with the *vegan* package. The number of distinct ARG in phylogroups or STs was referred to ARG richness.

## Funding

This work was partially supported by the “Fondation pour la Recherche Médicale” (Equipe FRM 2016, grant 325 number DEQ20161136698) and by the Direction Générale des Armées (project FastGeneII).

## Acknowledgements

We are grateful to Prof. Alexander Mellmann (University of Münster, Germany) for providing the *E. coli* strain containing the *Bacteroides uniformis* beta-lactamases encoding gene, and to James R. Johnson (University of Minnesota, MN), Lance B. Price (University of Washington, DC) and Maliha Aziz ((University of Washington, DC) for helping in understanding the *S. aureus/E. coli blaZ* finding. We are grateful to Olivier Clermont for his help in the database curation. Last, we thank Olivier Tenaillon for his proofreading of the manuscript.

## Notes

### Competing Interest Statement

The authors have declared no competing interest.

## References

Beghain J, Bridier-Nahmias A, Le Nagard H, Denamur E, Clermont O. 2018. ClermonTyping: an easy-to-use and accurate in silico method for *Escherichia* genus strain phylotyping. Microb Genomics 4.

Billard-Pomares T, Fouteau S, Jacquet ME, Roche D, Barbe V, Castellanos M, Bouet JY, Cruveiller S, Médigue C, Blanco J, et al. 2014. Characterization of a P1-Like Bacteriophage Carrying an SHV-2 Extended-Spectrum β-Lactamase from an *Escherichia coli* Strain. Antimicrob Agents Chemother 58: 6550–6557.

Bohlin J, Eldholm V, Pettersson JHO, Brynildsrud O, Snipen L. 2017. The nucleotide composition of microbial genomes indicates differential patterns of selection on core and accessory genomes. BMC Genomics 18: 151.

Bonnet R. 2004. Growing Group of Extended-Spectrum β-Lactamases: the CTX-M Enzymes. Antimicrob Agents Chemother48: 1–14.

Branger C, Ledda A, Billard-Pomares T, Doublet B, Barbe V, Roche D, Médigue C, Arlet G, Denamur E. 2019. Specialization of small non-conjugative plasmids in *Escherichia coli* according to their family types. Microb Genomics 5.

Branger C, Ledda A, Billard-Pomares T, Doublet B, Fouteau S, Barbe V, Roche D, Cruveiller S, Médigue C, Castellanos M, et al. 2018. Extended-spectrum β-lactamase-encoding genes are spreading on a wide range of *Escherichia coli* plasmids existing prior to the use of third-generation cephalosporins. Microb Genomics 4.

Branger C, Zamfir O, Geoffroy S, Laurans G, Arlet G, Thien HV, Gouriou S, Picard B, Denamur E. 2005. Genetic Background of *Escherichia coli* and Extended-spectrum β-Lactamase Type. Emerg Infect Dis 11: 54–61.

Buchfink B, Xie C, Huson DH. 2015. Fast and sensitive protein alignment using DIAMOND. Nat Methods 12: 59–60.

Carattoli A, Zankari E, García-Fernández A, Larsen MV, Lund O, Villa L, Aarestrup FM, Hasman H. 2014. In Silico Detection and Typing of Plasmids using PlasmidFinder and Plasmid Multilocus Sequence Typing. Antimicrob Agents Chemother 58: 3895–3903.

Clermont O, Dixit OVA, Vangchhia B, Condamine B, Dion S, Bridier-Nahmias A, Denamur E, Gordon D. 2019. Characterization and rapid identification of phylogroup G in *Escherichia coli*, a lineage with high virulence and antibiotic resistance potential. Environ Microbiol 21: 3107–3117.

Deng M, Zhu M-H, Li J-J, Bi S, Sheng Z-K, Hu F-S, Zhang J-J, Chen W, Xue X-W, Sheng J-F, et al. 2014. Molecular Epidemiology and Mechanisms of Tigecycline Resistance in Clinical Isolates of *Acinetobacter baumannii* from a Chinese University Hospital. Antimicrob Agents Chemother 58: 297–303.

Deschamps C, Clermont O, Hipeaux MC, Arlet G, Denamur E, Branger C. 2009. Multiple acquisitions of CTX-M plasmids in the rare D2 genotype of *Escherichia coli* provide evidence for convergent evolution. Microbiology, 155: 1656–1668.

Domingo J, Baeza-Centurion P, Lehner B. 2019. The Causes and Consequences of Genetic Interactions (Epistasis). Annu Rev Genomics Hum Genet 20: 433–460.

Escobar-Páramo P, Clermont O, Blanc-Potard A-B, Bui H, Le Bouguénec C, Denamur E. 2004. A Specific Genetic Background Is Required for Acquisition and Expression of Virulence Factors in *Escherichia coli*. Mol Biol Evol 21: 1085–1094.

Feldgarden M, Brover V, Haft DH, Prasad AB, Slotta DJ, Tolstoy I, Tyson GH, Zhao S, Hsu C-H, McDermott PF, et al. 2019. Using the NCBI AMRFinder Tool to Determine Antimicrobial Resistance Genotype-Phenotype Correlations Within a Collection of NARMS Isolates. bioRxiv 550707.

Fu L, Niu B, Zhu Z, Wu S, Li W. 2012. CD-HIT: accelerated for clustering the next-generation sequencing data. Bioinformatics 28: 3150–3152.

Gupta SK, Padmanabhan BR, Diene SM, Lopez-Rojas R, Kempf M, Landraud L, Rolain J-M. 2014. ARG-ANNOT, a New Bioinformatic Tool To Discover Antibiotic Resistance Genes in Bacterial Genomes. Antimicrob Agents Chemother 58: 212–220.

Jain R, Rivera MC, Lake JA. 1999. Horizontal gene transfer among genomes: The complexity hypothesis. Proc Natl Acad Sci 96: 3801–3806.

Jia B, Raphenya AR, Alcock B, Waglechner N, Guo P, Tsang KK, Lago BA, Dave BM, Pereira S, Sharma AN, et al. 2017. CARD 2017: expansion and model-centric curation of the comprehensive antibiotic resistance database. Nucleic Acids Res 45: D566–D573.

Joensen KG, Scheutz F, Lund O, Hasman H, Kaas RS, Nielsen EM, Aarestrup FM. 2014. Real-Time Whole-Genome Sequencing for Routine Typing, Surveillance, and Outbreak Detection of Verotoxigenic *Escherichia coli*. J Clin Microbiol 52: 1501–1510.

Johnson JR, Goullet P, Picard B, Moseley SL, Roberts PL, Stamm WE. 1991. Association of carboxylesterase B electrophoretic pattern with presence and expression of urovirulence factor determinants and antimicrobial resistance among strains of *Escherichia coli* that cause urosepsis. Infect Immun 59: 2311–2315.

Johnson JR, Orskov I, Orskov F, Goullet P, Picard B, Moseley SL, Roberts PL, Stamm WE. 1994. O, K, and H Antigens Predict Virulence Factors, Carboxylesterase B Pattern, Antimicrobial Resistance, and Host Compromise among *Escherichia coli* Strains Causing Urosepsis. J Infect Dis 169: 119–126.

Jolley KA, Maiden MC. 2010. BIGSdb: Scalable analysis of bacterial genome variation at the population level. BMC Bioinformatics 11: 595.

Kim S-W, Karns JS, Kessel JASV, Haley BJ. 2017. Genome Sequences of 30 *Escherichia coli* O157:H7 Isolates Recovered from a Single Dairy Farm and Its Associated Off-Site Heifer-Raising Facility. Genome Announc 5.

Kondratyeva K, Salmon-Divon M, Navon-Venezia S. 2020. Meta-analysis of Pandemic *Escherichia coli* ST131 Plasmidome Proves Restricted Plasmid-clade Associations. Sci Rep 10: 36.

Leski TA, Bangura U, Jimmy DH, Ansumana R, Lizewski SE, Stenger DA, Taitt CR, Vora GJ. 2013. Multidrug-resistant tet(X)-containing hospital isolates in Sierra Leone. Int J Antimicrob Agents 42: 83–86.

Li J, Jia H, Cai X, Zhong H, Feng Q, Sunagawa S, Arumugam M, Kultima JR, Prifti E, Nielsen T, et al. 2014. An integrated catalog of reference genes in the human gut microbiome. Nat Biotechnol 32: 834–841.

Li W, Godzik A. 2006. Cd-hit: a fast program for clustering and comparing large sets of protein or nucleotide sequences. Bioinformatics 22: 1658–1659.

Liu B, Pop M. 2009. ARDB--Antibiotic Resistance Genes Database. Nucleic Acids Res 37: D443–447.

Manges AR, Johnson JR, Foxman B, O’Bryan TT, Fullerton KE, Riley LW. 2001. Widespread Distribution of Urinary Tract Infections Caused by a Multidrug-Resistant *Escherichia coli* Clonal Group. N Engl J Med 345: 1007–1013.

Mellmann A, Bletz S, Böking T, Kipp F, Becker K, Schultes A, Prior K, Harmsen D. 2016. Real-Time Genome Sequencing of Resistant Bacteria Provides Precision Infection Control in an Institutional Setting. J Clin Microbiol 54: 2874–2881.

Munk P, Knudsen BE, Lukjancenko O, Duarte ASR, Van Gompel L, Luiken REC, Smit LAM, Schmitt H, Garcia AD, Hansen RB, et al. 2018. Abundance and diversity of the faecal resistome in slaughter pigs and broilers in nine European countries. Nat Microbiol 3: 898–908.

Nicolas-Chanoine M-H, Bertrand X, Madec J-Y. 2014. *Escherichia coli* ST131, an Intriguing Clonal Group. Clin Microbiol Rev 27: 543–574.

Ondov BD, Treangen TJ, Melsted P, Mallonee AB, Bergman NH, Koren S, Phillippy AM. 2016. Mash: fast genome and metagenome distance estimation using MinHash. Genome Biol 17: 132.

Porse A, Schou TS, Munck C, Ellabaan MMH, Sommer MOA. 2018. Biochemical mechanisms determine the functional compatibility of heterologous genes. Nat Commun 9: 522.

Ruppé E, Ghozlane A, Tap J, Pons N, Alvarez A-S, Maziers N, Cuesta T, Hernando-Amado S, Clares I, Martínez JL, et al. 2019. Prediction of the intestinal resistome by a three-dimensional structurebased method. Nat Microbiol 4: 112–123.

Sahl JW, Morris CR, Emberger J, Fraser CM, Ochieng JB, Juma J, Fields B, Breiman RF, Gilmour M, Nataro JP, et al. 2015. Defining the Phylogenomics of *Shigella* Species: a Pathway to Diagnostics. J Clin Microbiol 53: 951–960.

Slanetz LW, Bartley CH. 1957. Numbers of Enterococci in Water, Sewage, and Feces Determined by the Membrane Filter Technique with an Improved Medium. J Bacteriol 74: 591–595.

Smati M, Clermont O, Bleibtreu A, Fourreau F, David A, Daubié A-S, Hignard C, Loison O, Picard B, Denamur E. 2015. Quantitative analysis of commensal *Escherichia coli* populations reveals host-specific enterotypes at the intra-species level. MicrobiologyOpen 4: 604–615.

Sommer MOA, Dantas G, Church GM. 2009. Functional Characterization of the Antibiotic Resistance Reservoir in the Human Microflora. Science 325: 1128–1131.

Stalder T, Barraud O, Casellas M, Dagot C, Ploy M-C. 2012. Integron Involvement in Environmental Spread of Antibiotic Resistance. Front Microbiol 3.

Tenaillon O, Skurnik D, Picard B, Denamur E. 2010. The population genetics of commensal *Escherichia coli*. Nat Rev Microbiol 8: 207–217.

Wallace JC, Port JA, Smith MN, Faustman EM. 2017. FARME DB: a functional antibiotic resistance element database. Database 2017.

Zankari E, Hasman H, Cosentino S, Vestergaard M, Rasmussen S, Lund O, Aarestrup FM, Larsen MV. 2012. Identification of acquired antimicrobial resistance genes. J Antimicrob Chemother 67: 2640–2644.

Zhou Z, Alikhan N-F, Mohamed K, Fan Y, Group the AS, Achtman M, Brown D, Chattaway M, Dallman T, Delahay R, et al. 2020. The EnteroBase user’s guide, with case studies on *Salmonella* transmissions, *Yersinia pestis* phylogeny, and *Escherichia* core genomic diversity. Genome Res 30: 138–152.

