## Supplemental material for "Phylum barrier and *Escherichia coli* intra-species phylogeny drive the acquisition of resistome in *E. coli*"

| Beta-lactamase sub-family | 100 % identity | Variants | Frequency |
|---|---|---|---|
| $bla_{TEM}$ | 17,848 | 516 | 48.96 |
| $bla_{CTX\text{-}M}$ | 10,472 | 168 | 28.37 |
| $bla_{OXA\text{-}1\text{-}like}$ | 4,034 | 41 | 10.86 |
| $bla_{CMY\text{-}2\text{-}like}$ | 1,938 | 56 | 5.32 |
| $bla_{SHV}$ | 392 | 126 | 1.38 |
| $bla_{OXA\text{-}48\text{-}like}$ | 315 | 0 | 0.84 |
| $bla_{NDM}$ | 296 | 232 | 1.41 |
| $bla_{KPC}$ | 186 | 0 | 0.50 |
| $bla_{LAP}$ | 155 | 6 | 0.43 |
| $bla_{CARB}$ | 138 | 0 | 0.37 |
| $bla_{OXA\text{-}10\text{-}like}$ | 121 | 6 | 0.34 |
| $bla_{DHA}$ | 119 | 21 | 0.37 |
| $bla_{HER}$ | 102 | 0 | 0.27 |
| $bla_{OXA\text{-}2\text{-}like}$ | 77 | 0 | 0.21 |
| $bla_{OXA\text{-}PR\text{-}like}$ | 56 | 0 | 0.15 |
| $bla_{VIM\text{-}1}$ | 25 | 0 | 0.07 |
| $bla_{IMP}$ | 17 | 0 | 0.05 |
| $bla_{GES}$ | 12 | 0 | 0.03 |
| $bla_{VEB}$ | 9 | 0 | 0.02 |
| $bla_{SCO}$ | 8 | 0 | 0.02 |
| $bla_{ACC}$ | 6 | 1 | 0.02 |
| $bla_{SFO}$ | 6 | 0 | 0.02 |
| $bla_{OXA\text{-}5\text{-}like}$ | 2 | 0 | 0.01 |
| $bla_{FOX}$ | 1 | 0 | 0.00 |
| $bla_{MOX}$ | 0 | 1 | 0.00 |
| $bla_{ROB}$ | 0 | 1 | 0.00 |
| $bla_{SED}$ | 0 | 1 | 0.00 |

**Supplementary Table 1: The 27 sub-families of beta-lactamases found in *E. coli*.** 100% identity refers to strain containing exactly the sequence of the beta-lactamase and variants correspond to genes sharing at least 80% of identity and/or coverage with those from AMRFinder.

| Mustard Gene ID | AMRFinder Gene ID | Identity (%) | Coverage | Identity |
|---|---|---|---|---|
| MC3.MG14.AS1.GP1.C34954.G50 | 1028085905\|WP_063844403.1\|NG_047726\|1\|1\|dfrA3\|dfrA3\|trimethoprim-resistant_dihydrofolate_reductase_DfrA3 | 51.2 | 98.2 | 100 % identity |
| MC3.MG8.AS1.GP1.C7085.G8 | 1028085905\|WP_063844403.1\|NG_047726\|1\|1\|dfrA3\|dfrA3\|trimethoprim-resistant_dihydrofolate_reductase_DfrA3 | 52.5 | 98.2 | 100 % identity |
| MC3.MG117.AS1.GP1.C23735.G8 | 1028085975\|WP_063844473.1\|NG_047739\|1\|1\|dfrA7\|dfrA7\|trimethoprim-resistant_dihydrofolate_reductase_DfrA7 | 29.5 | 80.4 | 100 % identity |
| MC3.MG276.AS1.GP1.C9618.G2 | 1028110235\|WP_063864605.1\|NG_049991\|1\|1\|blaSHV-101\|blaSHV\|class_A_beta-lactamase_SHV-101 | 99.7 | 99.7 | 100 % identity |
| MC3.MG8.AS1.GP1.C15753.G2 | 1061158354\|WP_069174570.1\|NG_051497\|1\|1\|fosA6\|fosA_gen\|fosfomycin_resistance_glutathione_transferase_FosA6 | 99.3 | 99.3 | 100 % identity |
| MC3.MG304.AS1.GP1.C14695.G2 | 1105502238\|WP_071846388.1\|NG_052524\|1\|1\|dfrA5\|dfrA5\|trimethoprim-resistant_dihydrofolate_reductase_DfrA5 | 99.3 | 96.2 | 100 % identity |
| MC3.MG317.AS1.GP1.C48340.G4 | 1391852856\|WP_109545041.1\|NG_056002\|1\|1\|aph(3'')-Ib\|aph(3'')-Ib\|aminoglycoside_O-phosphotransferase_APH(3'')-Ib | 100.0 | 99.6 | 100 % identity |
| MC3.MG173.AS1.GP1.C14217.G2 | 445940466\|WP_000018321.1\|NG_056047\|1\|1\|aph(3')-Ia\|aph(3')-Ia\|aminoglycoside_O-phosphotransferase_APH(3')-Ia | 100.0 | 99.6 | 100 % identity |
| MC3.MG78.AS1.GP1.C21721.G2 | 445949202\|WP_000027057.1\|NG_050145\|1\|1\|blaTEM-1\|blaTEM\|class_A_broad-spectrum_beta-lactamase_TEM-1 | 100.0 | 99.7 | 100 % identity |
| MC3.MG276.AS1.GP1.C7768.G6 | 445953162\|WP_000031017.1\|NG_047988\|1\|1\|mph(B)\|mph(B)\|Mph(B)_family_macrolide_2'-phosphotransferase | 100.0 | 99.7 | 100 % identity |
| MC3.MG209.AS1.GP1.C20136.G1 | 445956565\|WP_000034420.1\|NG_048120\|1\|1\|sul3\|sul3\|sulfonamide-resistant_dihydropteroate_synthase_Sul3 | 35.1 | 97.7 | 100 % identity |
| MC3.MG276.AS1.GP1.C1921.G2 | 446181177\|WP_000259032.1\|NG_048081\|1\|1\|sul1\|sul1\|sulfonamide-resistant_dihydropteroate_synthase_Sul1 | 99.6 | 99.6 | 100 % identity |
| MC3.MG2.AS1.GP1.C12473.G2 | 446403113\|WP_000480968.1\|NG_047464\|1\|1\|aph(6)-Id\|aph(6)-Id\|aminoglycoside_O-phosphotransferase_APH(6)-Id | 100.0 | 99.6 | 100 % identity |
| MC3.MG377.AS1.GP1.C83819.G1 | 446479600\|WP_000557454.1\|NG_047251\|1\|1\|aac(3)-IId\|aac(3)-II\|aminoglycoside_N-acetyltransferase_AAC(3)-IId | 100.0 | 99.7 | 100 % identity |
| MC3.MG250.AS1.GP1.C34776.G2 | 446493211\|WP_000571065.1\|NG_047741\|1\|1\|dfrA8\|dfrA8\|trimethoprim-resistant_dihydrofolate_reductase_DfrA8 | 100.0 | 99.4 | 100 % identity |
| MC3.MG276.AS1.GP1.C7508.G1 | 446700209\|WP_000777555.1\|NG_047678\|1\|1\|dfrA1\|dfrA1\|trimethoprim-resistant_dihydrofolate_reductase_DfrA1 | 100.0 | 96.2 | 100 % identity |
| MC3.MG192.AS1.GP1.C13240.G2 | 447129060\|WP_001206316.1\|NG_047324\|1\|1\|aadA1\|aadA1\|ANT(3'')-Ia_family_aminoglycoside_nucleotidyltransferase_AadA1 | 100.0 | 99.6 | 100 % identity |
| MC3.MG96.AS1.GP1.C8327.G4 | 487657633\|WP_001749985.1\|NG_047723\|1\|1\|dfrA27\|dfrA\|trimethoprim-resistant_dihydrofolate_reductase_DfrA27 | 33.3 | 80.4 | 100 % identity |
| MC3.MG68.AS1.GP1.C34414.G1 | 488221018\|WP_002292226.1\|NG_047794\|1\|1\|erm(B)\|erm(B)\|23S_rRNA_(adenine(2058)-N(6))-methyltransferase_Erm(B) | 100.0 | 99.6 | 100 % identity |
| MC3.MG149.AS1.GP1.C12456.G1 | 488642079\|WP_002578722.1\|NG_047322\|1\|1\|aad9\|aad9\| | 100.0 | 99.6 | 100 % identity |

| | | | | |
|---|---|---|---|---|
| | ANT(9)_family_aminoglycoside_nucleotidyltransferase | | | |
| MC3.MG60.AS1.GP1.C4251.G1 | 489068003\|WP_002977989.1\|NG_048758\|1\|1\|blaCGA-1\|blaCGA\|class_A_extended-spectrum_beta-lactamase_CGA-1 | 37.2 | 99.7 | 100 % identity |
| MC3.MG241.AS1.GP1.C60081.G1 | 499192666\|WP_010890206.1\|NG_048106\|1\|1\|sul2\|sul2\|sulfonamide-resistant_dihydropteroate_synthase_Sul2 | 99.6 | 99.6 | 100 % identity |
| MC3.MG243.AS1.GP1.C43768.G2 | 499446086\|WP_011133550.1\|NG_047776\|1\|1\|erm(33)\|erm(33)\|23S_rRNA_(adenine(2058)-N(6))-methyltransferase_Erm(33) | 27.8 | 97.5 | 100 % identity |
| MC3.MG14.AS1.GP1.C34954.G45 | 499601687\|WP_011282421.1\|NG_047815\|1\|1\|erm(C)\|erm(C)\|23S_rRNA_(adenine(2058)-N(6))-methyltransferase_Erm(C) | 28.6 | 69.4 | 100 % identity |
| MC3.MG14.AS1.GP1.C28907.G43 | 504873498\|WP_015060600.1\|NG_048308\|1\|1\|tet(X)\|tet(X)\|tetracycline-inactivating_monooxygenase_Tet(X) | 24.3 | 80.7 | 100 % identity |
| MC3.MG257.AS1.GP1.C42691.G1 | 545212893\|WP_021529707.1\|NG_047696\|1\|1\|dfrA14\|dfrA14\|trimethoprim-resistant_dihydrofolate_reductase_DfrA14 | 100.0 | 99.4 | 100 % identity |
| MC3.MG327.AS1.GP1.C13288.G1 | 695270536\|WP_032492605.1\|NG_047891\|1\|1\|fosC2\|fosC2\|FosC2_family_fosfomycin_resistance_glutathione_transferase | 28.6 | 78.9 | 100 % identity |
| MC3.MG342.AS1.GP1.C30231.G1 | 917745009\|WP_052259183.1\|NG_047784\|1\|1\|erm(42)\|erm(42)\|23S_rRNA_(adenine(2058)-N(6))-methyltransferase_Erm(42) | 46.1 | 92.1 | 100 % identity |
| MC3.MG8.AS1.GP1.C3157.G1 | | | | 100 % identity |
| MC3.MG382.AS1.GP1.C47625.G1 | | | | 100 % identity |
| MC3.MG278.AS1.GP1.C23765.G6 | | | | 100 % identity |
| MC3.MG276.AS1.GP1.C9721.G1 | | | | 100 % identity |
| MC3.MG14.AS1.GP1.C5528.G2 | | | | 100 % identity |
| MC3.MG14.AS1.GP1.C34954.G4 | | | | 100 % identity |
| MC3.MG14.AS1.GP1.C23888.G23 | | | | 100 % identity |
| MC3.MG14.AS1.GP1.C1185.G20 | | | | 100 % identity |
| MC3.MG117.AS1.GP1.C16515.G24 | | | | 100 % identity |
| MC3.MG257.AS1.GP1.C45437.G3 | 488758799\|WP_002682030.1\|NG_047825\|1\|1\|erm(F)\|erm(F)\|23S_rRNA_(adenine(2058)-N(6))-methyltransferase_Erm(F) | 89.8 | 99.6 | Variant |
| MC3.MG338.AS1.GP1.C31454.G4 | 446926004\|WP_001003260.1\|NG_055988\|1\|1\|erm(C)\|erm(C)\|23S_rRNA_(adenine(2058)-N(6))-methyltransferase_Erm(C) | 28.2 | 94.3 | Variant |
| MC3.MG169.AS1.GP1.C31521.G2 | 1028099858\|WP_063856112.1\|NG_048219\|1\|1\|tet(M)\|tet(M)\|tetracycline_resistance_ribosomal_protection_protein_Tet(M) | 98.6 | 99.8 | Variant |
| MC3.MG153.AS1.GP1.C20144.G2 | 949781494\|WP_057098416.1\|NG_047408\|1\|1\|aph(2")-Ih\|aph(2")-Ih\|aminoglycoside_O-phosphotransferase_APH(2")-Ih | 25.0 | 82.6 | Variant |
| MC3.MG338.AS1.GP1.C21977.G2 | 504873498\|WP_015060600.1\|NG_048308\|1\|1\|tet(X)\|tet(X)\|tetracycline-inactivating_monooxygenase_Tet(X) | 23.3 | 82.5 | Variant |
| MC3.MG170.AS1.GP1. | | | | Variant |

| | | | | |
|---|---|---|---|---|
| C17246.G1 | | | | |
| MC3.MG229.AS1.GP1.C3473.G14 | | | | Variant |
| MC3.MG170.AS1.GP1.C7561.G2 | | | | Variant |
| MC3.MG170.AS1.GP1.C1586.G1 | 1391852845\|WP_109545032.1\|NG_050471\|1\|1\|qnrB11\|qnrB\|quinolone_resistance_pentapeptide_repeat_protein_QnrB11 | 99.1 | 54.4 | Variant |
| MC3.MG338.AS1.GP1.C13378.G11 | 740629697\|WP_038415208.1\|NG_050405\|1\|1\|fosA\|fosA-154989\|fosfomycin_resistance_glutathione_transferase_FosA | 95.7 | 99.3 | Variant |
| MC3.MG229.AS1.GP1.C35745.G8 | 1028085894\|WP_063844392.1\|NG_047724\|1\|1\|dfrA28\|dfrA\|trimethoprim-resistant_dihydrofolate_reductase_DfrA28 | 32.6 | 80.4 | Variant |
| MC3.MG3.AS1.GP1.C19448.G6 | | | | Variant |
| MC3.MG229.AS1.GP1.C13544.G33 | 445956565\|WP_000034420.1\|NG_048120\|1\|1\|sul3\|sul3\|sulfonamide-resistant_dihydropteroate_synthase_Sul3 | 35.5 | 97.7 | Variant |
| MC3.MG338.AS1.GP1.C20089.G2 | | | | Variant |
| MC3.MG377.AS1.GP1.C45339.G1 | | | | Variant |
| MC3.MG229.AS1.GP1.C7137.G6 | | | | Variant |
| MC3.MG8.AS1.GP1.C7085.G5 | 446926004\|WP_001003260.1\|NG_055988\|1\|1\|erm(C)\|erm(C)\|23S_rRNA_(adenine(2058)-N(6))-methyltransferase_Erm(C) | 28.6 | 69.4 | Variant |
| MC3.MG338.AS1.GP1.C6822.G6 | | | | Variant |
| MC3.MG222.AS1.GP1.C20581.G1 | | | | Variant |
| MC3.MG41.AS1.GP1.C9847.G2 | 495926503\|WP_008651082.1\|NG_048305\|1\|1\|tet(X)\|tet(X)\|tetracycline-inactivating_monooxygenase_Tet(X) | 100.0 | 99.7 | Variant |
| MC3.MG118.AS1.GP1.C40284.G3 | | | | Variant |
| MC3.MG338.AS1.GP1.C10074.G1 | | | | Variant |
| MC3.MG8.AS1.GP1.C7085.G8 | 1028085905\|WP_063844403.1\|NG_047726\|1\|1\|dfrA3\|dfrA3\|trimethoprim-resistant_dihydrofolate_reductase_DfrA3 | 52.5 | 98.2 | Variant |

**Supplementary Table 2: Genes of Mustard database found in *E. coli*.** “100% identity” refers to strain containing exactly the sequence of the genes and “variant” corresponds to variants of the sequence with at least 80% of identity/coverage.

| ResfinderFG Gene ID | AMRFinder Gene ID | Identity (%) | Coverage | Identity |
|---|---|---|---|---|
| AAC\|GQ343136.1\|feces\|GEN | 446479598\|WP_000557452.1\|NG_047244\|1\|1\|aac(3)-IIa\|aac(3)-IIa\|aminoglycoside_N-acetyltransferase_AAC(3)-IIa | 100.0 | 97.9 | 100 % identity |
| AAC\|GQ343185.1\|feces\|SIS | 1028080938\|WP_063840321.1\|NG_051711\|1\|1\|aac(6')-Ib-cr5\|aac(6')-Ib-cr\|fluoroquinolone-acetylating_aminoglycoside_6'-N-acetyltransferase_AAC(6')-Ib-cr5 | 100.0 | 99.5 | 100 % identity |
| AAC\|KU544318.1\|sewage\|GEN | 504199701\|WP_014386803.1\|NG_047998\|1\|1\|mph(G)\|mph(G)\|Mph(G)_family_macrolide_2'-phosphotransferase | 62.4 | 69.5 | 100 % identity |
| AAC\|KU544324.1\|sewage\|GEN | 446479600\|WP_000557454.1\|NG_047251\|1\|1\|aac(3)-IId\|aac(3)-II\|aminoglycoside_N-acetyltransferase_AAC(3)-IId | 100.0 | 99.7 | 100 % identity |
| ANT\|KU544724.1\|sewage\|GEN | 446303947\|WP_000381802.1\|NG_047387\|1\|1\|ant(2")-Ia\|ant(2")-Ia\|aminoglycoside_nucleotidyltransferase_ANT(2")-Ia | 100.0 | 99.4 | 100 % identity |
| beta_lactamase\|GQ343005.1\|feces\|AMX | 446161735\|WP_000239590.1\|NG_048935\|1\|1\|blaCTX-M-15\|blaCTX-M\|class_A_extended-spectrum_beta-lactamase_CTX-M-15 | 100.0 | 99.7 | 100 % identity |
| beta_lactamase\|KF626746.1\|feces\|PEN | | | | 100 % identity |
| beta_lactamase\|KU543694.1\|feces\|AMX | 445949202\|WP_000027057.1\|NG_050145\|1\|1\|blaTEM-1\|blaTEM\|class_A_broad-spectrum_beta-lactamase_TEM-1 | 100.0 | 99.7 | 100 % identity |
| beta_lactamase\|KU543700.1\|feces\|CTX | | | | 100 % identity |
| beta_lactamase\|KU543892.1\|feces\|PEN | 445949195\|WP_000027050.1\|NG_050162\|1\|1\|blaTEM-116\|blaTEM\|class_A_broad-spectrum_beta-lactamase_TEM-116 | 100.0 | 99.7 | 100 % identity |
| beta_lactamase\|KU543933.1\|sewage\|AMX | 490302154\|WP_004197546.1\|NG_049978\|1\|1\|blaSCO-1\|blaSCO\|class_A_beta-lactamase_SCO-1 | 100.0 | 99.7 | 100 % identity |
| beta_lactamase\|KU543938.1\|sewage\|AMX | 446629385\|WP_000706731.1\|NG_050317\|1\|1\|blaVEB-1\|blaVEB\|class_A_extended-spectrum_beta-lactamase_VEB-1 | 100.0 | 99.7 | 100 % identity |
| beta_lactamase\|KU543962.1\|sewage\|AMX | 485701606\|WP_001334766.1\|NG_049392\|1\|1\|blaOXA-1\|blaOXA-1_like\|oxacillin-hydrolyzing_class_D_beta-lactamase_OXA-1 | 100.0 | 99.6 | 100 % identity |
| beta_lactamase\|KU543973.1\|sewage\|AMX | 501858675\|WP_012658785.1\|NG_049137\|1\|1\|blaGES-5\|blaGES\|carbapenem-hydrolyzing_class_A_beta-lactamase_GES-5 | 100.0 | 99.7 | 100 % identity |
| beta_lactamase\|KU543983.1\|sewage\|AMX | 1175118064\|WP_081666691.1\|NG_056482\|1\|1\|blaLHK-1\|blaLHK\|class_C_beta-lactamase_LHK-1 | 44.1 | 88.7 | 100 % identity |
| beta_lactamase\|KU544013.1\|sewage\|ATM | 486436156\|WP_001617865.1\|NG_048929\|1\|1\|blaCTX-M-14\|blaCTX-M\|class_A_extended-spectrum_beta-lactamase_CTX-M-14 | 99.6 | 96.9 | 100 % identity |
| beta_lactamase\|KU544434.1\|sewage\|PIP | 446769134\|WP_000846390.1\|NG_049393\|1\|1\|blaOXA-10\|blaOXA-10_like\|oxacillin-hydrolyzing_class_D_beta-lactamase_OXA-10 | 100.0 | 99.6 | 100 % identity |
| beta_lactamase\|KU544887.1\|feces\|CAZ | 1028104515\|WP_063859836.1\|NG_048829\|1\|1\|blaCMY-34\|blaCMY\|class_C_beta-lactamase_CMY-34 | 99.7 | 99.7 | 100 % identity |
| beta_lactamase\|KU545465.1\|feces\|PEN | 1175118064\|WP_081666691.1\|NG_056482\|1\|1\|blaLHK-1\|blaLHK\|class_C_beta-lactamase_LHK-1 | 43.8 | 88.7 | 100 % identity |
| beta_lactamase\|KU545603.1\|feces\|PEN | | | | 100 % identity |
| beta_lactamase\|KU545796.1\|feces\|AMX | 742671073\|WP_038976851.1\|NG_050215\|1\|1\|blaTEM-176\|blaTEM\|class_A_beta-lactamase_TEM- | 100.0 | 99.7 | 100 % identity |

| | | | | |
|---|---|---|---|---|
| | 176 | | | |
| beta_lactamase\|KU546246.1\|feces\|AMX | 490280343\|WP_004176269.1\|NG_050000\|1\|1\|blaSHV-11\|blaSHV\|class_A_broad-spectrum_beta-lactamase_SHV-11 | 100.0 | 99.7 | 100 % identity |
| beta_lactamase\|KU546296.1\|feces\|CAZ | 502953499\|WP_013188475.1\|NG_049016\|1\|1\|blaCTX-M-65\|blaCTX-M\|class_A_extended-spectrum_beta-lactamase_CTX-M-65 | 100.0 | 96.9 | 100 % identity |
| beta_lactamase\|KU546394.1\|feces\|AMX | 488993270\|WP_002904004.1\|NG_050590\|1\|1\|blaSHV-12\|blaSHV\|class_A_extended-spectrum_beta-lactamase_SHV-12 | 100.0 | 99.7 | 100 % identity |
| beta_lactamase\|KU546399.1\|feces\|AMX | | | | 100 % identity |
| beta_lactamase\|KU546551.1\|feces\|AMX | 487858008\|WP_001931474.1\|NG_048722\|1\|1\|blaCARB-2\|blaPSE\|PSE_family_carbenicillin-hydrolyzing_class_A_beta-lactamase_CARB-2 | 100.0 | 99.7 | 100 % identity |
| beta_lactamase\|KU547585.1\|feces\|AMX | 1028104535\|WP_063859856.1\|NG_048859\|1\|1\|blaCMY-63\|blaCMY\|class_C_beta-lactamase_CMY-63 | 75.8 | 99.5 | 100 % identity |
| beta_lactamase\|KU548170.1\|latrine\|PEN | 490360692\|WP_004240455.1\|NG_049058\|1\|1\|blaDHA-13\|blaDHA\|class_C_beta-lactamase_DHA-13 | 99.6 | 69.7 | 100 % identity |
| beta_lactamase\|KU606101.1\|feces\|PEN | | | | 100 % identity |
| beta_lactamase\|KU606137.1\|feces\|PIP | | | | 100 % identity |
| beta_lactamase\|KU606886.1\|feces\|CAZ | | | | 100 % identity |
| beta_lactamase\|KU607183.1\|feces\|CAZ | | | | 100 % identity |
| cat\|KU544029.1\|sewage\|CHL | 446108382\|WP_000186237.1\|NG_047604\|1\|1\|catB3\|catB3\|type_B-3_chloramphenicol_O-acetyltransferase_CatB3 | 100.0 | 99.5 | 100 % identity |
| dfr\|KF630308.1\|feces\|SXT | 447006469\|WP_001083725.1\|NG_047689\|1\|1\|dfrA12\|dfrA12\|trimethoprim-resistant_dihydrofolate_reductase_DfrA12 | 100.0 | 99.4 | 100 % identity |
| efflux_pump\|KU546555.1\|feces\|CHL | 446136267\|WP_000214122.1\|NG_047877\|1\|1\|floR\|floR\|chloramphenicol/florfenicol_efflux_MFS_transporter_FloR | 99.8 | 99.8 | 100 % identity |
| qnr\|KU548326.1\|latrine\|CIP | 1028111843\|WP_063866054.1\|NG_050498\|1\|1\|qnrB36\|qnrB\|quinolone_resistance_pentapeptide_repeat_protein_QnrB36 | 99.5 | 99.5 | 100 % identity |
| tet_efflux\|KU543718.1\|feces\|TET | 447011812\|WP_001089068.1\|NG_048163\|1\|1\|tet(B)\|tet(B)\|tetracycline_efflux_MFS_transporter_Tet(B) | 100.0 | 99.8 | 100 % identity |
| tet_efflux\|KU544792.1\|sewage\|TET | 485651500\|WP_001297013.1\|NG_048177\|1\|1\|tet(C)\|tet(C)\|tetracycline_efflux_MFS_transporter_Tet(C) | 100.0 | 99.7 | 100 % identity |
| tet_efflux\|KU545892.1\|feces\|TET | 447011816\|WP_001089072.1\|NG_048161\|1\|1\|tet(B)\|tet(B)\|tetracycline_efflux_MFS_transporter_Tet(B) | 100.0 | 99.8 | 100 % identity |
| tet_efflux\|KU547117.1\|latrine\|TET | 665836803\|WP_031204920.1\|NG_048123\|1\|1\|tet(31)\|tet(31)\|tetracycline_efflux_MFS_transporter_Tet(31) | 100.0 | 99.8 | 100 % identity |
| tet_efflux\|KU548366.1\|latrine\|TET | 446764192\|WP_000841448.1\|NG_048175\|1\|1\|tet(C)\|tet(C)\|tetracycline_efflux_MFS_transporter_Tet(C) | 100.0 | 99.7 | 100 % identity |
| tet_protection\|KU545260.1\|feces\|TET | 446614381\|WP_000691727.1\|NG_048250\|1\|1\|tet(M)\|tet(M)\|tetracycline_resistance_ribosomal_protection_protein_Tet(M) | 100.0 | 99.8 | 100 % identity |

| | | | | |
|---|---|---|---|---|
| tetX\|KU547176.1\|latrine\|TET | 1028100309\|WP_063856444.1\|NG_048306\|1\|1\|tet(X)\|tet(X)\|tetracycline-inactivating_monooxygenase_Tet(X) | 96.0 | 99.7 | 100 % identity |
| tetX\|KU547431.1\|feces\|TET | 504873815\|WP_015060917.1\|NG_048307\|1\|1\|tet(X)\|tet(X)\|tetracycline-inactivating_monooxygenase_Tet(X) | 100.0 | 99.7 | 100 % identity |
| 16S_rRNA_methyltransferase\|KU548774.1\|feces\|GEN | 557680565\|WP_023434793.1\|NG_051537\|1\|1\|rmtF2\|rmtF\|16S_rRNA_(guanine(1405)-N(7))-methyltransferase_RmtF2 | 98.8 | 99.6 | Variant |
| beta_lactamase\|GQ343075.1\|feces\|CDR | 765425998\|WP_044699195.1\|NG_048843\|1\|1\|blaCMY-48\|blaCMY\|class_C_beta-lactamase_CMY-48 | 99.7 | 99.7 | Variant |
| beta_lactamase\|KF629077.1\|feces\|PEN | | | | Variant |
| beta_lactamase\|KF630336.1\|feces\|PEN | 1028104684\|WP_063859951.1\|NG_048924\|1\|1\|blaCTX-M-132\|blaCTX-M\|class_A_extended-spectrum_beta-lactamase_CTX-M-132 | 83.5 | 99.7 | Variant |
| beta_lactamase\|KU543927.1\|sewage\|AMX | 446930417\|WP_001007673.1\|NG_049496\|1\|1\|blaOXA-2\|blaOXA-2_like\|oxacillin-hydrolyzing_class_D_beta-lactamase_OXA-2 | 100.0 | 99.6 | Variant |
| beta_lactamase\|KU544007.1\|sewage\|AMX | 446930417\|WP_001007673.1\|NG_049496\|1\|1\|blaOXA-2\|blaOXA-2_like\|oxacillin-hydrolyzing_class_D_beta-lactamase_OXA-2 | 99.6 | 99.6 | Variant |
| beta_lactamase\|KU544162.1\|sewage\|CTX | 503015905\|WP_013250881.1\|NG_049111\|1\|1\|blaGES-1\|blaGES\|class_A_extended-spectrum_beta-lactamase_GES-1 | 100.0 | 99.7 | Variant |
| beta_lactamase\|KU544353.1\|sewage\|PEN | 446769134\|WP_000846390.1\|NG_049393\|1\|1\|blaOXA-10\|blaOXA-10_like\|oxacillin-hydrolyzing_class_D_beta-lactamase_OXA-10 | 99.6 | 99.6 | Variant |
| beta_lactamase\|KU544486.1\|sewage\|AMX | 695269134\|WP_032491311.1\|NG_049688\|1\|1\|blaOXA-4\|blaOXA-1_like\|OXA-1_family_oxacillin-hydrolyzing_class_D_beta-lactamase_OXA-4 | 99.6 | 99.6 | Variant |
| beta_lactamase\|KU545125.1\|feces\|AMX | 445949205\|WP_000027060.1\|NG_050218\|1\|1\|blaTEM-181\|blaTEM\|class_A_beta-lactamase_TEM-181 | 99.7 | 99.7 | Variant |
| beta_lactamase\|KU545264.1\|feces\|AMX | 1028104338\|WP_063859734.1\|NG_048790\|1\|1\|blaCMY-119\|blaCMY\|class_C_beta-lactamase_CMY-119 | 99.7 | 99.7 | Variant |
| beta_lactamase\|KU545266.1\|feces\|AMX | | | | Variant |
| beta_lactamase\|KU545271.1\|feces\|ATM | | | | Variant |
| beta_lactamase\|KU545287.1\|feces\|TZP | | | | Variant |
| beta_lactamase\|KU545493.1\|feces\|AMX | | | | Variant |
| beta_lactamase\|KU546390.1\|feces\|AMX | | | | Variant |
| beta_lactamase\|KU547393.1\|feces\|AMX | | | | Variant |
| beta_lactamase\|KU547568.1\|feces\|PEN | | | | Variant |
| beta_lactamase\|KU548068.1\|latrine\|PIP | 695270053\|WP_032492122.1\|NG_049428\|1\|1\|blaOXA-129\|blaOXA-5_like\|OXA-5_family_class_D_beta-lactamase_OXA-129 | 94.0 | 99.6 | Variant |
| beta_lactamase\|KU548547.1\|soil\|AMX | 446930417\|WP_001007673.1\|NG_049496\|1\|1\|blaOXA-2\|blaOXA-2_like\|oxacillin-hydrolyzing_class_D_beta-lactamase_OXA-2 | 99.3 | 99.6 | Variant |

| | | | | |
|---|---|---|---|---|
| beta_lactamase\|KU548878.1\|feces\|AMX | 1028105818\|WP_063860843.1\|NG_049312\|1\|1\|blaMOX-10\|blaMOX\|CMY-1/MOX_family_class_C_beta-lactamase_MOX-10 | 45.5 | 91.1 | Variant |
| beta_lactamase\|KU606564.1\|feces\|AMP | | | | Variant |
| beta_lactamase\|KU607207.1\|feces\|PIP | | | | Variant |
| beta_lactamase\|KU607800.1\|feces\|PEN | 1028104330\|WP_063859728.1\|NG_048789\|1\|1\|blaCMY-118\|blaCMY\|class_C_beta-lactamase_CMY-118 | 92.6 | 99.5 | Variant |
| beta_lactamase\|KU607811.1\|feces\|PEN | | | | Variant |
| beta_lactamase\|KU608079.1\|feces\|AMP | | | | Variant |
| cat\|KU547990.1\|latrine\|CHL | 498321738\|WP_010635894.1\|NG_052738\|1\|1\|catB8\|catB8\|type_B-3_chloramphenicol_O-acetyltransferase_CatB8 | 100.0 | 99.5 | Variant |
| efflux_pump\|KU547150.1\|latrine\|CHL | 447179520\|WP_001256776.1\|NG_047647\|1\|1\|cmlA1\|cmlA1\|chloramphenicol_efflux_MFS_transporter_CmlA1 | 99.8 | 99.8 | Variant |
| efflux_pump\|KU548456.1\|soil\|CHL | 487652940\|WP_001747811.1\|NG_047861\|1\|1\|floR\|floR\|chloramphenicol/florfenicol_efflux_MFS_transporter_FloR | 99.8 | 99.8 | Variant |
| efflux_pump\|KU548715.1\|soil\|TET | 447179517\|WP_001256773.1\|NG_047655\|1\|1\|cmlA6\|cmlA\|chloramphenicol_efflux_MFS_transporter_CmlA6 | 100.0 | 99.8 | Variant |
| tet_efflux\|GQ343144.1\|feces\|MIN | 1028099444\|WP_063855887.1\|NG_048151\|1\|1\|tet(A)\|tet(A)\|tetracycline_efflux_MFS_transporter_Tet(A) | 99.6 | 61.8 | Variant |
| tet_efflux\|GQ343149.1\|feces\|OXY | 1028099445\|WP_063855888.1\|NG_048152\|1\|1\|tet(A)\|tet(A)\|tetracycline_efflux_MFS_transporter_Tet(A) | 99.7 | 99.8 | Variant |
| tet_efflux\|KU544451.1\|sewage\|TET | 446764190\|WP_000841446.1\|NG_048181\|1\|1\|tet(C)\|tet(C)\|tetracycline_efflux_MFS_transporter_Tet(C) | 99.5 | 99.7 | Variant |
| tet_efflux\|KU544465.1\|sewage\|TGC | 500229186\|WP_011899270.1\|NG_048188\|1\|1\|tet(E)\|tet(E)\|tetracycline_efflux_MFS_transporter_Tet(E) | 99.8 | 99.8 | Variant |
| tet_efflux\|KU544473.1\|sewage\|TGC | 1028099571\|WP_063855982.1\|NG_048155\|1\|1\|tet(A)\|tet(A)\|tetracycline_efflux_MFS_transporter_Tet(A) | 99.5 | 99.8 | Variant |
| tet_efflux\|KU544912.1\|feces\|TET | 446726751\|WP_000804064.1\|NG_048154\|1\|1\|tet(A)\|tet(A)\|tetracycline_efflux_MFS_transporter_Tet(A) | 99.7 | 99.8 | Variant |
| tet_efflux\|KU545776.1\|feces\|TET | 487900204\|WP_001973670.1\|NG_048156\|1\|1\|tet(A)\|tet(A)\|tetracycline_efflux_MFS_transporter_Tet(A) | 100.0 | 99.8 | Variant |
| tet_efflux\|KU545910.1\|feces\|TET | 447180584\|WP_001257840.1\|NG_051907\|1\|1\|tet(G)\|tet(G)\|tetracycline_efflux_MFS_transporter_Tet(G) | 99.7 | 99.7 | Variant |
| tet_efflux\|KU546496.1\|feces\|TGC | 1028099442\|WP_063855886.1\|NG_048150\|1\|1\|tet(A)\|tet(A)\|tetracycline_efflux_MFS_transporter_Tet(A) | 98.2 | 99.7 | Variant |
| tet_efflux\|KU548943.1\|feces\|TGC | 446726751\|WP_000804064.1\|NG_048154\|1\|1\|tet(A)\|tet(A)\|tetracycline_efflux_MFS_transporter_Tet(A) | 99.7 | 99.8 | Variant |
| tet_protection\|KU543820.1\|feces\|TET | 758866324\|WP_043029018.1\|NG_048244\|1\|1\|tet(M)\|tet(M)\|tetracycline_resistance_ribosomal_protection_protein_Tet(M) | 99.5 | 99.8 | Variant |
| tet_protection\|KU545044.1\|feces\|TET | 912687134\|WP_050232631.1\|NG_048246\|1\|1\|tet(M)\|tet(M)\|tetracycline_resistance_ribosomal_protection_protein_Tet(M) | 99.1 | 99.8 | Variant |

| | | | | |
|---|---|---|---|---|
| tet_protection\|KU545174.1\|feces\|TET | 501919555\|WP_012669445.1\|NG_048264\|1\|1\|tet(O)\|tet(O)\|tetracycline_resistance_ribosomal_protection_protein_Tet(O) | 99.8 | 99.8 | Variant |
| tet_protection\|KU546373.1\|feces\|TET | 1028100163\|WP_063856393.1\|NG_048229\|1\|1\|tet(M)\|tet(M)\|tetracycline_resistance_ribosomal_protection_protein_Tet(M) | 99.2 | 99.8 | Variant |
| tet_protection\|KU547111.1\|latrine\|TET | 1028100163\|WP_063856393.1\|NG_048229\|1\|1\|tet(M)\|tet(M)\|tetracycline_resistance_ribosomal_protection_protein_Tet(M) | 98.9 | 99.8 | Variant |
| tet_protection\|KU548962.1\|feces\|TET | 1028100163\|WP_063856393.1\|NG_048229\|1\|1\|tet(M)\|tet(M)\|tetracycline_resistance_ribosomal_protection_protein_Tet(M) | 98.8 | 87.7 | Variant |
| tetX\|KU547125.1\|latrine\|TET | 1028100309\|WP_063856444.1\|NG_048306\|1\|1\|tet(X)\|tet(X)\|tetracycline-inactivating_monooxygenase_Tet(X) | 98.7 | 99.7 | Variant |
| tetX\|KU547753.1\|latrine\|TET | 495926503\|WP_008651082.1\|NG_048305\|1\|1\|tet(X)\|tet(X)\|tetracycline-inactivating_monooxygenase_Tet(X) | 99.7 | 98.5 | Variant |
| van_ligase\|GQ343123.1\|feces\|CYC | 1028100561\|WP_063856696.1\|NG_048370\|1\|1\|vanG2\|vanG2\|D-alanine--D-serine_ligase_VanG2 | 39.0 | 99.7 | Variant |
| van_ligase\|GQ343119.1\|feces\|CYC | 1028100510\|WP_063856645.1\|NG_048345\|1\|1\|vanC1\|vanC1\|D-alanine--D-serine_ligase_VanC1 | 34.7 | 98.0 | Variant |
| efflux_pump\|KU546654.1\|feces\|CHL | 446017870\|WP_000095725.1\|NG_047648\|1\|1\|cmlA1\|cmlA1\|chloramphenicol_efflux_MFS_transporter_CmlA1 | 100.0 | 99.8 | Variant |
| efflux_pump\|KU544501.1\|sewage\|CHL | 447179520\|WP_001256776.1\|NG_047647\|1\|1\|cmlA1\|cmlA1\|chloramphenicol_efflux_MFS_transporter_CmlA1 | 100.0 | 99.8 | Variant |
| efflux_pump\|KU544040.1\|sewage\|CHL | 501257754\|WP_012300772.1\|NG_051436\|1\|1\|cmlA5\|cmlA5\|chloramphenicol_efflux_MFS_transporter_CmlA5 | 100.0 | 99.8 | Variant |
| beta_lactamase\|KU607183.1\|feces\|CAZ | | | | Variant |
| beta_lactamase\|KU606886.1\|feces\|CAZ | | | | Variant |
| beta_lactamase\|KU606137.1\|feces\|PIP | | | | Variant |
| beta_lactamase\|KU606101.1\|feces\|PEN | | | | Variant |
| beta_lactamase\|KU545603.1\|feces\|PEN | | | | Variant |
| beta_lactamase\|KU543700.1\|feces\|CTX | | | | Variant |
| beta_lactamase\|KF626746.1\|feces\|PEN | | | | Variant |
| AAC\|GQ343134.1\|feces\|GEN | 446479598\|WP_000557452.1\|NG_047244\|1\|1\|aac(3)-IIa\|aac(3)-IIa\|aminoglycoside_N-acetyltransferase_AAC(3)-IIa | 100.0 | 99.7 | Variant |

**Supplementary Table 3: Genes of ResFinderFG database found in *E. coli*.** “100% identity” refers to strain containing exactly the sequence of the genes and “variant” corresponds to variants of the sequence with at least 80% of identity/coverage.

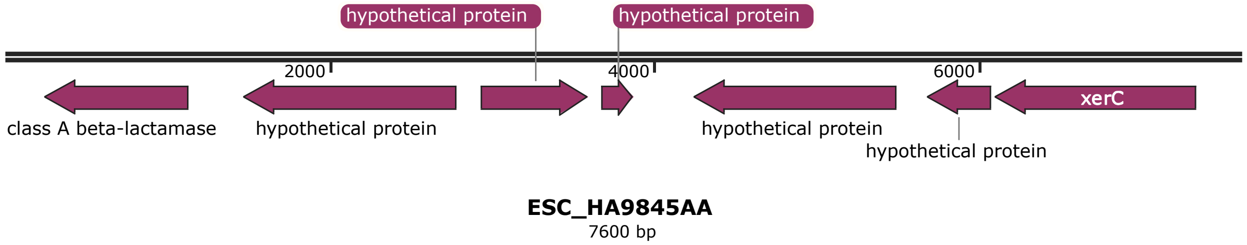


**Supplementary Figure 1: Description of the genes shared by the 7,600 bp contig of the strain matching against a *Bacteroides uniformis*.** *B. uniformis* genes are coloured in purple.

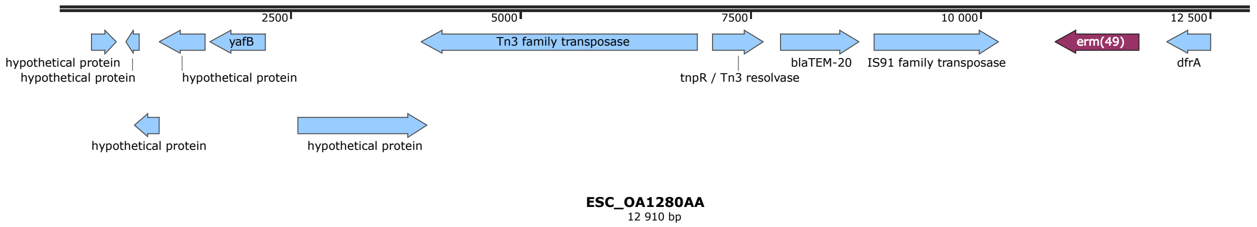


**Supplementary Figure 2: Description of the genes shared by the 12,910 bp contig of the strain matching against a *Bifidobacterium breve*.** *E. coli* genes are coloured in blue and *B. breve* genes in purple (71 bp in 3' of *erm*(49) also matching with *B. breve* genome).

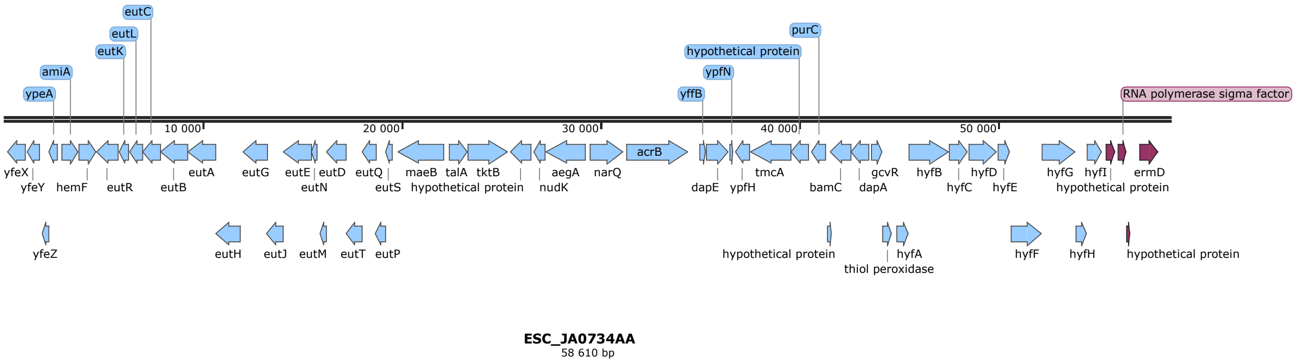


**Supplementary Figure 3: Description of the genes shared by the 58,610 bp contig of the strain matching against a *Clostridioides difficile*.** *E. coli* genes are coloured in blue and those from *C. difficile* in purple (25 bp in 5' of the first hypothetical protein also matching with *C. difficile).*

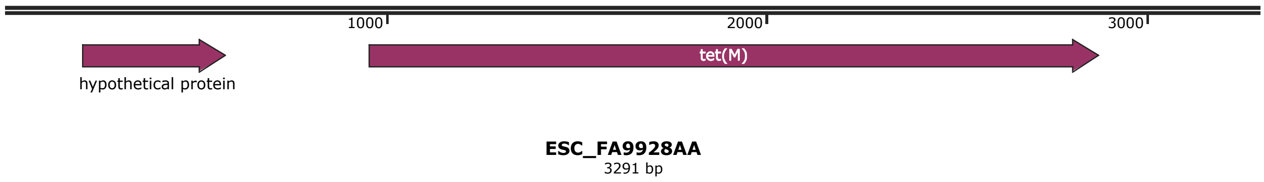


**Supplementary Figure 4: Description of the genes shared by the 3,291 bp contig of the strain matching against a *Clostridioides difficile*.** C*. difficile* genes are coloured in purple.
